# Focal adhesion membrane is dotted with protein islands and partitioned for molecular hop diffusion

**DOI:** 10.1101/2021.10.26.465868

**Authors:** Takahiro K. Fujiwara, Shinji Takeuchi, Ziya Kalay, Yosuke Nagai, Taka A. Tsunoyama, Thomas Kalkbrenner, Limin H. Chen, Akihiro C.E. Shibata, Kokoro Iwasawa, Ken P. Ritchie, Kenichi G.N. Suzuki, Akihiro Kusumi

## Abstract

Using the ultrafast camera system and new theories for hop diffusion described in the companion paper, we for the first time demonstrated that membrane molecules undergo hop diffusion among the compartments in the bulk *basal* plasma membrane (PM), with virtually the same compartment sizes (108 nm) as those in the bulk apical PM and the same dwell lifetimes within a compartment (10 and 24 ms for the phospholipid and transferrin receptor, respectively), suggesting that the basic structures and molecular dynamics are very similar in the bulk regions of the apical and basal PMs. Ultrafast PALM and single-molecule imaging revealed that the focal adhesion (FA) is mostly a fluid membrane, partitioned into ∼74-nm compartments where transferrin receptor and β3 integrin undergo hop diffusion, and that the FA membrane is sparsely dotted with 51-nm diameter paxillin islands, where many other FA proteins probably assemble (compartmentalized archipelago model). β3 integrin intermittently associates with the paxillin islands, dynamically linking them to the extracellular matrix.

**Summary:** An ultrafast camera with single fluorescent-molecule sensitivities developed by Fujiwara et al. reveals that the focal adhesion membrane is dotted with protein islands and partitioned for molecular hop diffusion, and integrin β3 molecules become temporarily immobilized at the islands.

## Introduction

In the companion paper (Fujiwara et al., 2021), we report the development of an ultrahigh-speed camera system that has enabled the fastest single fluorescent-molecule imaging ever performed. We achieved a 100-µs resolution with a 20-nm localization precision for single Cy3 molecules for a frame size of 14 x 14 µm^2^ (256 x 256 pixels), and a 33-µs resolution with a 34-nm localization precision for a frame size of 7.1 x 6.2 µm^2^ (128 x 112 pixels) (**Table 1** in the companion paper). This is probably the ultimate rate possible with the currently available fluorescent probes, due to their limited throughputs.

**Table 1.**
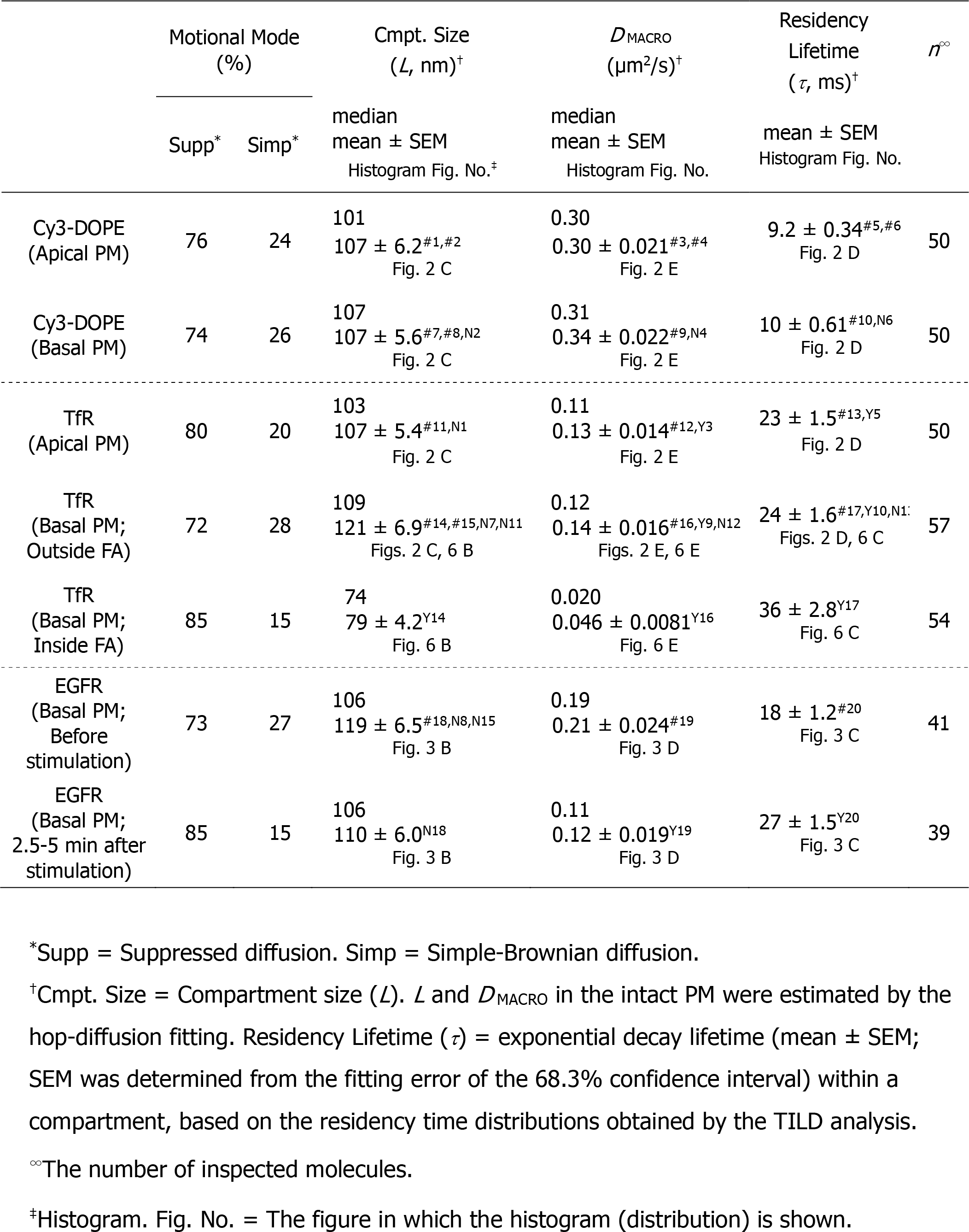
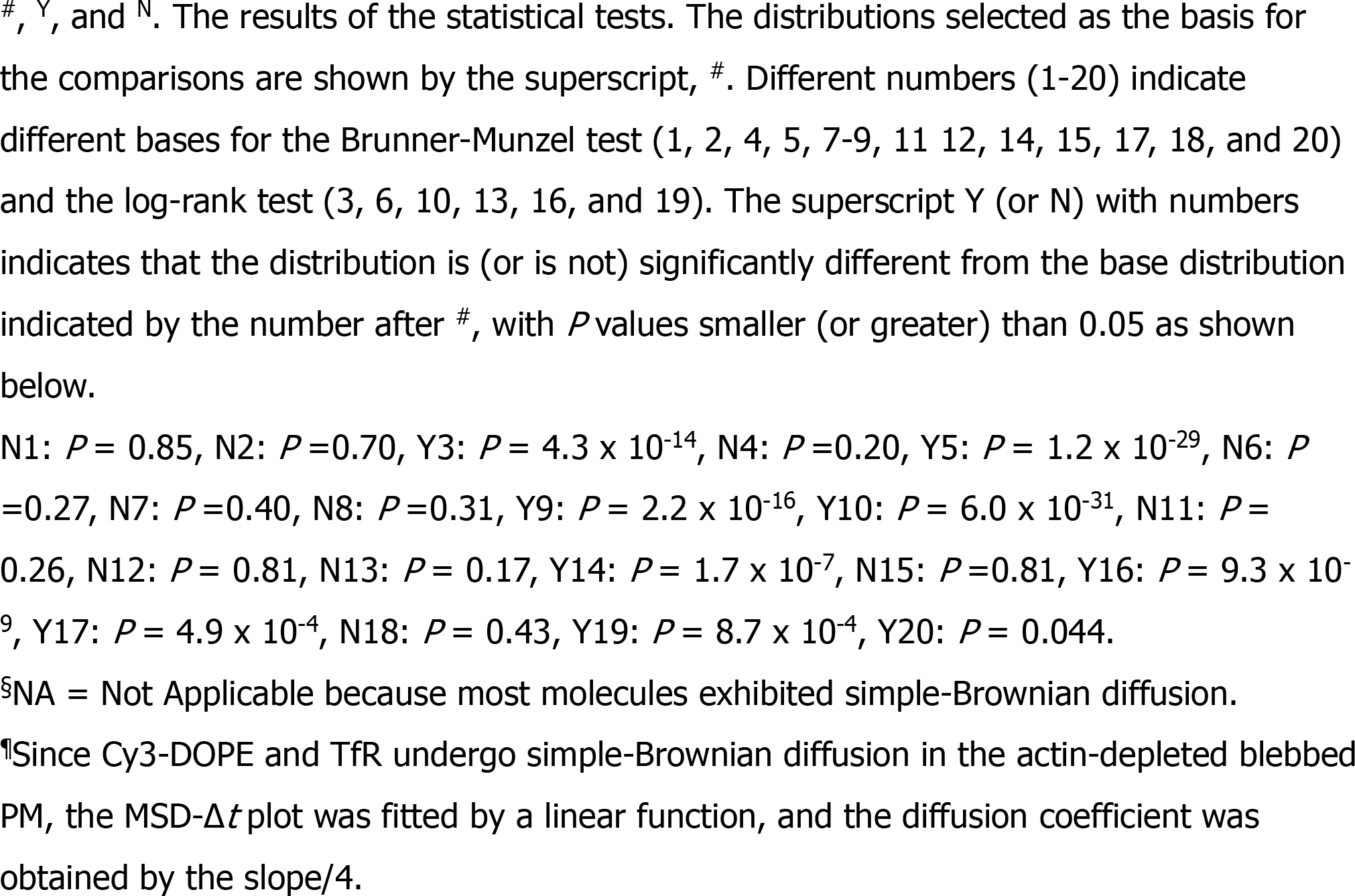
Motional mode, compartment size (*L*), *D* _MACRO_, and dwell lifetime (*τ*), characterizing the hop diffusions of Cy3-DOPE, TfR, and EGFR in apical and basal PMs of T24 cells.

Previously, hop diffusion of the phospholipids and membrane proteins in the PM was detectable only by single-particle tracking using *40-nm diameter-gold probes* and high-speed bright-field microscopy, at a camera frame rate of 40 kHz (a time resolution of 25 µs; Fujiwara et al., 2002, 2016; Murase et al., 2004). To obtain reasonable single-particle localization precisions at this frame rate for tracking the fast dynamics of single molecules in the PM, large colloidal gold probes and bright-field microscopy were necessary. Our original intention when developing the ultrafast camera for single *fluorescent molecule* imaging was to make such fast single-particle observations possible with fluorescent probes, which are more prevalent and favorable than 40-nm gold probes.

Therefore, in the companion paper, the applicability of the developed camera system to cell biology was examined by testing whether it can detect the hop diffusion of membrane molecules in the PM. It nicely reproduced the ultrafast gold-particle tracking data, using a fluorescently-labeled phospholipid and transferrin receptor (TfR), a transmembrane protein.

Using this ultrafast camera system and the theories developed for analyzing the movements of molecules undergoing hop diffusion in the compartmentalized plasma membrane (PM) (**Fig. S5** in the companion paper), we were, for the first time, able to measure the distributions of the residency times within a compartment. These distributions could be represented by a single exponential function, as predicted by the developed theory (the decay time constant provides the dwell lifetime of the molecule within a compartment). The use of fluorescent probes allowed us to perform a diffusion anomaly analysis in a 5 order of magnitude time range (from 0.044 ms to 2 s). These new results further supported the picket-fence model, based on the actin-based membrane skeleton meshes (fences) and transmembrane picket proteins anchored to and aligned along the fences (**Fig. 1, A** and **B**; Fujiwara et al., 2002; Morone et al., 2006).

**Figure 1.**
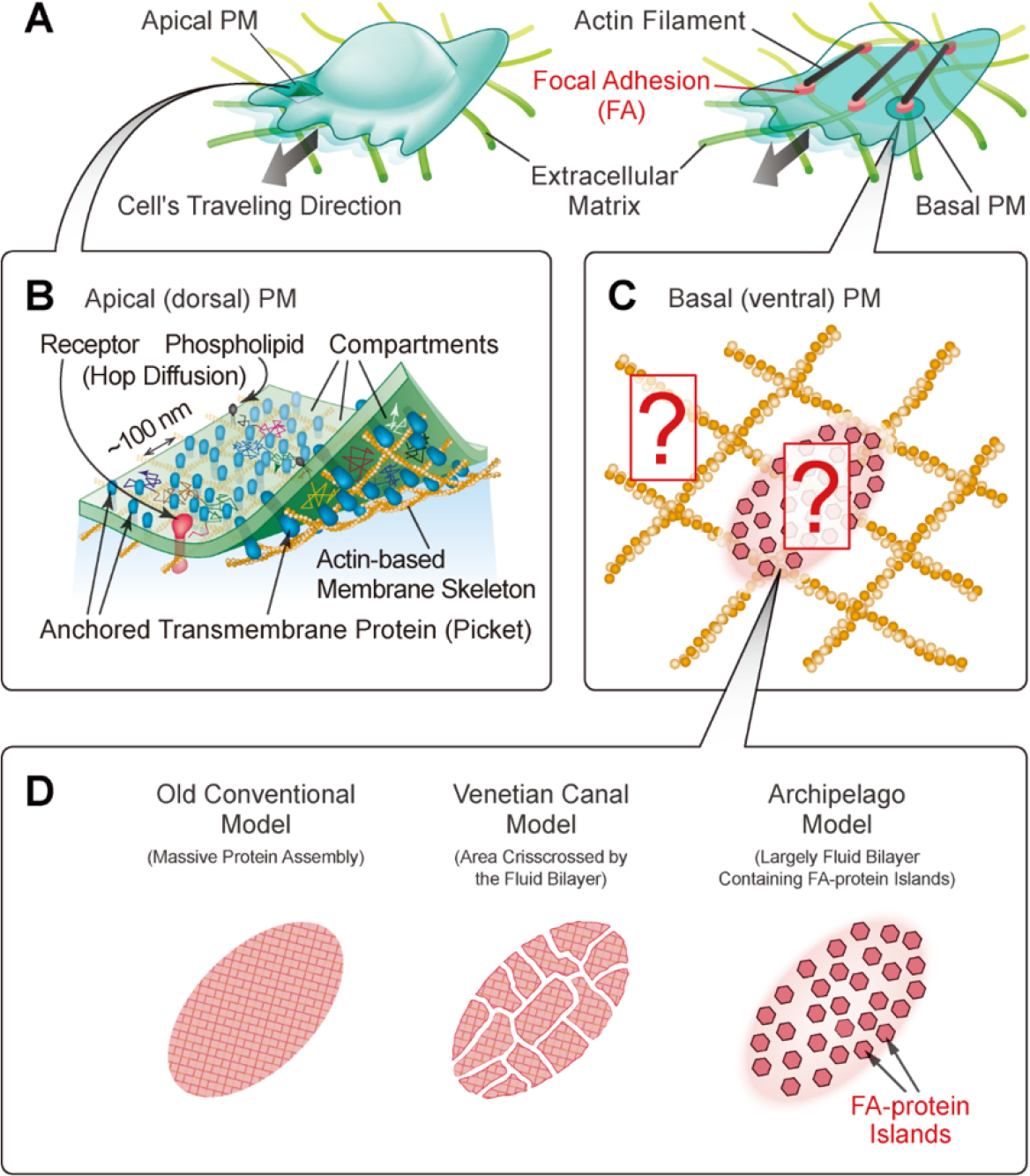
The purposes of the study: ultrahigh-speed single molecule dynamics in the basal PM including the FA region, and elucidation of how the FA fits into the view of the actin-induced compartmentalization. (**A**) The apical (dorsal) PM (**left**) and the basal (ventral) PM (**right**) compared in this research. The basal PM contains the FAs (red ellipsoids), which structurally and mechanically link the intracellular actin filaments (black) to the extracellular matrix (green filaments), providing key scaffolds for cell migration in/on the extracellular matrix. (**B**) The apical PM was previously found to be compartmentalized by the actin-based membrane-skeleton meshes (fences; brown mesh) and rows of transmembrane-protein pickets anchored to and aligned along the actin fence (blue molecules), which induce the hop diffusion of virtually all membrane molecules in the apical PM. (**C**) Two major questions about the basal PM. (1) Do membrane molecules undergo hop diffusion in the *basal* (ventral) PM (is it compartmentalized)? If yes, what are the compartment sizes and the dwell lifetimes of membrane molecules within a compartment? Are they similar to those found in the apical PM? (2) Are the FAs compartmentalized? (**D**) Three schematic FA models proposed so far. See the main text.

However, since the gold probes cannot enter the space between the *basal (ventral) PM* and the coverslip, and cannot label molecules in the basal PM, all of the gold-particle data and the test results for ultrafast single fluorescent-molecule imaging were obtained in the *apical (dorsal) PM*. With the development of the ultrafast camera system to image single fluorescent molecules, we can now, for the first time, experimentally address whether the molecular dynamics and compartmentalization in the basal PM are similar to those in the apical PM (questions outside the FA region in **Fig. 1** **C**). Using fluorescent probes, we found that both the phospholipid and transmembrane proteins, TfR and EGF receptor (EGFR), undergo hop diffusion in the *basal* PM, and that the compartment sizes and the dwell lifetimes of the diffusing molecules within a compartment in the basal PM are virtually the same as those in the apical PM (**Fig. 1, C and D**).

We examined a specialized membrane domain in the basal PM, the focal adhesion (FA). The FA is a large micron-scale structure in the PM that serves as a scaffold for cellular attachment to and migration in/on the extracellular matrix (**Fig. 1 A**; Nayal et al., 2006; Shroff et al., 2008; Gardel et al., 2010; Xia and Kanchanawong, 2017; Yamada and Sixt, 2019). Integrins are receptors in the FA that bind to various extracellular matrix proteins and link them to the actin filaments in the cytoplasm, by way of other FA cytoplasmic proteins (Parsons et al., 2010; Humphries et al., 2019).

Using single fluorescent-molecule tracking at slower rates, we previously demonstrated that non-FA proteins, such as Thy1 and transferrin receptor (TfR), readily enter the FA zone, diffuse there, and exit again to the bulk PM (Shibata et al., 2012). Integrin molecules also freely enter the FA zone by diffusion and exit again to the bulk PM, but often undergo temporary immobilizations while inside the FA zone (Shibata et al., 2012; Rossier et al., 2012; Tsunoyama et al., 2018; Orré et al., 2021). The diffusion coefficients in the FA region were smaller only by factors of 2 - 3 as compared with those in the bulk PM. Such high mobility within the FA zone suggested that the FA domain largely consists of *fluid membrane*. Previously, the FA was often considered to be a micron-scale, massive assembly of various proteins (like a large continent in the bulk fluid PM) or that subdivided by “canals” crisscrossing the land, like the city plan of Venice (Venetian canal model) (**Fig. 1 D**, **left and middle**). However, according to these models, the macroscopic diffusion of membrane molecules would be virtually blocked or considerably slowed in the FA (by an order of magnitude or more in the macroscopic diffusion coefficient), despite the presence of the fluid membrane (canals) (Holcman et al., 2011; Saxton, 1982, 2010). Therefore, these models are incongruent with the results of single-molecule dynamics studies of membrane molecules in the FA (Shibata et al., 2012; Rossier et al., 2012; Tsunoyama et al., 2018; Orré et al., 2021).

Furthermore, previous PALM and dSTORM observations of paxillin, a representative FA structural protein, revealed the presence of paxillin clusters scattered in the FA region (Shroff et al., 2008; Changede and Sheetz, 2017; Deschout et al., 2017; Orré et al., 2021). These images by themselves do not exclude the massive continent and Venetian city map models. However, considering that the FA region largely consists of the fluid membrane, we proposed that paxillin is predominantly distributed as clusters in the fluid membrane within the FA (Shibata et al., 2012, 2013). Paxillin reportedly interacts extensively with FA proteins such as talin, vinculin, FAK, kindlin, ILK, and integrins, which in turn bind to each other and/or other FA proteins, including RIAM, VASP, and actinin, as well as actin and/or PI(4,5)P_2_ for the PM anchorage (Liu et al., 2000; Krause et al., 2003; Sjöblom et al., 2008; Calderwood et al., 2013; Sulzmaier et al., 2014; Yang et al., 2014; Bays et al., 2017; López-Colomé et al., 2017). Therefore, we proposed that the paxillin clusters would contain many other FA proteins, and thus we refer to them as “FA-protein islands” in the sense of “extended paxillin islands” and the FA architecture of the fluid membrane sea dotted with the FA-protein islands as the “FA archipelago model” (**Fig. 1 D** **right**) (Shibata et al., 2012, 2013; also see Patla et al., 2010; Rossier et al., 2012; Levet et al., 2015; Changede and Sheetz, 2017; Spiess et al., 2018; Tsunoyama et al., 2018; Orré et al., 2021). The molecular compositions of the FA-protein islands would vary from island to island and also in time (Stutchbury et al., 2017), including the stoichiometry of the molecules in each island, since some islands might not even contain paxillin. Meanwhile, along the z axis (in 3D), the FA-protein islands would consist of layers of proteins with distinct molecular compositions (Kanchanawong et al., 2010; Liu et al., 2015). In the present report, we will use “paxillin islands” to refer to the direct experimental PALM results using mEos3.2-paxillin, and “FA-protein islands” to indicate the extended paxillin islands.

The critically important point with the archipelago model is that the FA zone is predominantly composed of fluid membrane (Shibata et al., 2012; Rossier et al., 2012). Therefore, in the present research, we focused more specifically on the properties and structures of the fluid membrane part of the FA, rather than the FA-protein islands, in the FA archipelago model (question inside the FA in **Fig. 1C**). Since the initial stages of the present research demonstrated that the basal PM is compartmentalized, we hoped to clarify whether the liquid membrane domain within the FA is also compartmentalized (**Fig. 1** **C**). We indeed found that this is the case, but the compartment size was smaller (74 nm vs. 109 nm in the bulk basal PM).

The developed ultrafast camera system described in the companion paper allows live-cell PALM observations with 0.33-10 s time resolutions for image sizes of 35.3 x 35.3 µm^2^ (640 x 640 pixels for data acquisition; 1 ms/frame acquisition x 333 – 10,000 frames with a 29-nm localization precision for mEos3.2). In this report, we describe a microscope that allowed us to simultaneously perform live-cell PALM for visualizing the paxillin islands and single fluorescent-molecule imaging for observing the movement of single β3-integrin molecules. The results showed that more than two-thirds of β3-integrin’s temporary immobilization events occurred at the paxillin islands.

## Results

### TfR and Cy3-DOPE undergo hop diffusion in the basal PM, exhibiting virtually the same compartment sizes and dwell lifetimes as those in the apical PM

Single molecules of TfR (TfR’s N-terminus fused to the Halo-tag protein, labeled with TMR) were observed in the bulk basal PM outside the FA region marked with mGFP-paxillin, at a frame rate of 6 kHz (0.167-ms resolution; 200 times faster than normal video rate) (**Fig. 2** **A top**; see **Movie S1 top**; throughout this work, human epithelial T24 cells were employed and observed at 37°C). The single-molecule trajectories can be conveniently analyzed by plotting the mean-square displacement (MSD) against the time interval (*Δt*), called the MSD-*Δt* plot (see **Fig. 4** **A** of the companion paper for further details).

**Figure 2.**
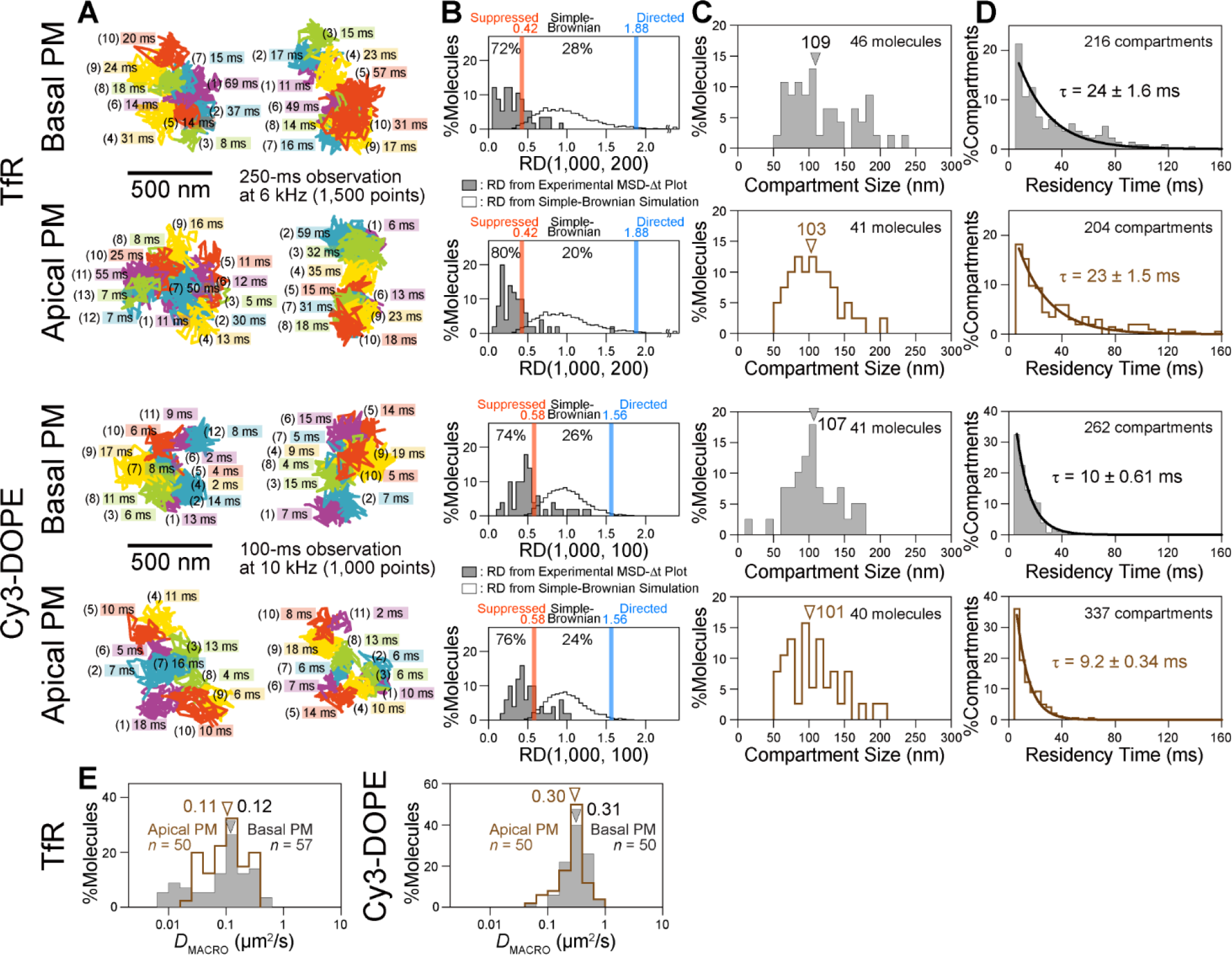
Ultrafast single fluorescent-molecule imaging of TfR and Cy3-DOPE revealed that the basal PM outside the FAs is compartmentalized like the apical PM and that the dwell lifetimes of TfR and Cy3-DOPE within a compartment in the basal PM outside the FAs are the same as those in the apical PM. (**A**) Typical ultrafast single fluorescent-molecule trajectories of TfR (6 kHz) and Cy3-DOPE (10 kHz), diffusing in the basal PM outside the FA region (**top**) and in the apical PM (**bottom**). The order of the compartments that the molecule visited (parenthesized integers) and the dwell lifetimes there, as determined by the TILD analysis (**Fig. S5** of the companion paper), are shown in the figure. TMR was used to label the Halo-tag protein fused to the cytoplasmic domain of TfR (N-terminus). Since Cy3 (and Alexa555), which exhibited the best throughput among all the dyes tested (**Figs. S2 and S3** in the companion paper), is membrane impermeable, the availability of a membrane-permeable dye, TMR, for ultrafast observations is very useful. The apical-PM data are reproduced from those in the companion paper (**Figs. 5** **B -D and 6 A**), except for the typical trajectories shown here. (**B**) Motional mode classification based on a parameter called <u>R</u>elative <u>D</u>eviation from ideal Brownian diffusion, *𝑅𝐷* (Fig. 5 **B** in the companion paper). Simple-Brownian particles exhibit the distributions of *𝑅𝐷* values (5,000 simulated particles) shown in the open histograms, and the red and blue vertical lines indicate the 2.5 percentiles from both ends of the distribution (*𝑅𝐷_min_* and *𝑅𝐷_MAX_*, respectively). The trajectories exhibiting *𝑅𝐷* values smaller than *𝑅𝐷_min_* are categorized into the suppressed diffusion mode. For details, see Fig. 5 **B** of the companion paper (also see Kusumi et al., 1993; Fujiwara et al., 2002; Murase et al., 2004; Suzuki et al., 2005). Shaded bars show the experimental *𝑅𝐷* distributions, and the percentages of molecules categorized into the suppressed and simple-Brownian diffusion modes are indicated. (**C**) Distributions of the compartment sizes determined from the hop-diffusion fitting of the MSD-Δ*t* plot for each TfR and Cy3-DOPE molecule. The MSD-Δ*t* plot data that could not be fit (due to large amounts of noise and the closeness of *D*_micro_ and *D*_MACRO_) were not included. Arrowheads indicate the median values. (**D**) Distributions of the TfR and Cy3-DOPE residency times within a compartment determined by the TILD analysis, with the best-fit single exponential functions (see the theory in **Fig. S5** in the companion paper). The decay time constants of these curves provide the dwell lifetimes. (**E**) Distribution of *D*_MACRO_ determined by the hop-diffusion fitting of the MSD-Δ*t* plot for each TfR and Cy3-DOPE molecule (shaded histogram, basal PM; open histogram, apical PM). Arrowheads indicate the median values. The statistical test methods, parameters (number of experiments), and *P* values throughout this report are summarized in **Table 1**.

Using the MSD-*Δt* plot for each TfR trajectory, the majorities of the trajectories were found to be categorized into the suppressed diffusion mode in the basal PM, like those in the apical PM (72 and 80%, respectively; **Fig. 2** **B** and **Table 1**; for the details of the classification method, see **Fig. 4** **B** of the companion paper). Consistently, the MSD-*Δt* plots of 81 and 82% of the trajectories obtained in the basal and apical PM, respectively, could be fitted by the equation describing the hop diffusion model (see the caption to **Fig. S5** in the companion paper), indicating that most TfR molecules undergo hop diffusion in the basal PM as well as in the apical PM. Interestingly, the median compartment size in the basal PM (109 nm) was quite comparable to that in the apical PM (103 nm) (**Fig. 2** **C**; non-significant difference; all statistical parameters and test results for diffusion parameters are summarized in **Table 1**).

The residency time of a TfR molecule within each compartment; i.e., the duration between two consecutive hop events, could be directly determined in each trajectory by using the analysis of <u>T</u>ransient <u>I</u>ncrease of the effective <u>L</u>ocal <u>D</u>iffusion (TILD analysis, see the companion paper), and the distributions of the TfR residency times within a compartment in the basal and apical PMs were obtained. Unexpectedly, the distribution in the basal PM appears very similar to that in the apical PM (**Fig. 2 D**). The distributions could be fitted by single exponential decay functions, as predicted by the hop diffusion theory (see the caption to **Fig. S5** in the companion paper), and the decay time constants provided the “dwell lifetimes” of TfR in a compartment of 24 ± 1.6 ms in the basal PM and 23 ± 1.5 ms in the apical PM (**Fig. 2 D**; from here on, we refer to the decay time constant as the dwell lifetime, whereas the dwell time of each molecule in each compartment is called the residency time). Namely, not only the compartment sizes, but also the TfR dwell lifetimes within a compartment were very similar to each other.

This was surprising, and so, the average residency time (*τ*) within a compartment was evaluated by another method. Namely, we used the equation *τ* = *L*^2^/4*D*_MACRO_, where *L* is the spacing of the 2D square lattice and *D*_MACRO_ is the long-term diffusion coefficient determined by the hop-diffusion fitting of the MSD-*Δt* plot (see the caption to **Fig. S5** of the companion paper). The median values of *L* were 109 and 103 nm (**Fig. 2** **C**) and those of *D*_MACRO_ were 0.12 and 0.11 µm^2^/s (**Fig. 2** **E**) in the basal and apical PMs, respectively. These provided the average residency times of 25 ms in the basal PM and 24 ms in the apical PM, in agreement with the values obtained by the TILD method. Namely, we conclude that the TfR’s dwell lifetimes in a compartment were virtually the same in both the basal and apical PMs.

The phospholipid, Cy3-labeled L-α-dioleoylphosphatidylethanolamine (Cy3-DOPE), also undergoes hop diffusion, exhibiting practically the same median compartment sizes as that for TfR in the basal PM and those for Cy3-DOPE and TfR in the apical PM (**Fig. 2** **C**). The dwell lifetime of Cy3-DOPE within a compartment was shorter than that of TfR by a factor of ∼2.5 in both the basal and apical PMs (**Fig. 2** **D**). Consistently, the *D*_MACRO_ values of Cy3-DOPE were the same in the basal and apical PMs (**Fig. 2** **E**). Overall, to our surprise, the bulk basal PM is compartmentalized in virtually the same manner as the apical PM, with regard to the compartment sizes and temporary confinements of TfR and Cy3-DOPE. Equality of the compartment size for TfR and Cy3-DOPE in the same membrane suggests that the underlying mechanisms for the compartmentalization are the same for these two molecules in both basal and apical PMs, consistent with the model of actin-based fences and transmembrane picket proteins anchored to and aligned along the fences.

Since the compartment sizes are so similar, the following possibility was examined: the hop diffusion might simply be an apparent phenomenon caused by the photo response non-uniformity (PRNU) of the developed camera system. PRNU might affect the single-molecule localization precisions through pixel-to-pixel variations in the sensitivity, which might make the molecules appear like those undergoing hop diffusion. The test result, shown in **Fig. S1**, indicated that PRNU of the developed camera is quite similar to that of EM-CCD camera, and would scarcely affects the single-molecule localization precisions.

Furthermore, as described later, we found different compartment sizes in the FA region, indicating that PRNU would not induce apparent hop-like movements for membrane molecules in the PM.

### Prolonged confinement of activated EGF receptor (EGFR) within a compartment

EGFR exists as both monomers and dimers before EGF stimulation, and they interconvert rapidly with a dimer lifetime of ∼13 s (Chung et al., 2010). After stimulation, EGFR tends to form rather stable dimers and greater oligomers for self-phosphorylation and activation (Chung et al., 2010; Low-Nam et al., 2011; Huang et al., 2016). We examined whether the EGF ligation of the receptor affects its lateral diffusion, because this will determine the rate of activated EGFR spreading along the PM by diffusion, which will provide a basis for a variety of mechanisms for EGFR signal propagation (Verveer et al., 2000; Reynolds et al., 2003; Koseska et al., 2020).

EGFR (fused to the Halo-tag protein at the cytoplasmic C-terminus of EGFR, labeled with TMR) was observed in the basal PM at the level of single molecules at a frame rate of 6 kHz (**Movie S2**). Due to the presence of non-labeled endogenous EGFR, even the fluorescent spots with the intensity of a single EGFR might represent EGFR dimers and greater oligomers. Virtually all of the fluorescent EGFR spots underwent hop diffusion both before and after ligation (**Fig. 3, A and B** and **Table 1**). Interestingly, their median compartment sizes were the same before and after stimulation (106 nm), and quite comparable to the compartment sizes in the basal PM found by observing TfR (109 nm) and DOPE (107 nm) (non-significant differences; **Table 1**), supporting the PM compartmentalization for all PM-associated molecules.

**Figure 3.**
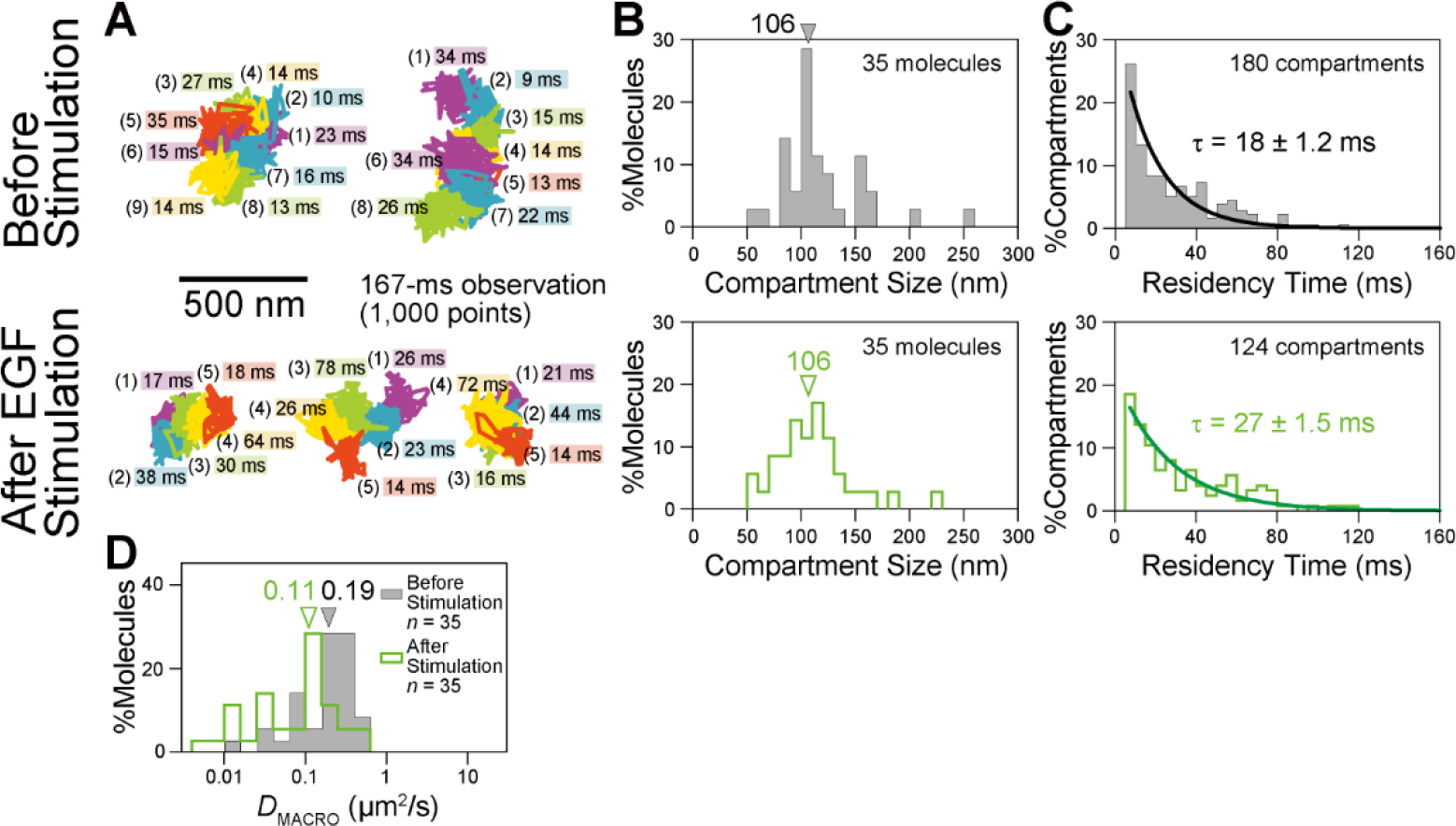
EGFR and ligand-engaged EGFR in the basal PM detected virtually the same compartment sizes as those found with TfR and Cy3-DOPE in the bulk basal PM, supporting the PM compartmentalization, and the dwell lifetime of the engaged EGFR was longer than that of non-engaged EGFR (same compartment sizes). After stimulation with 10 nM EGF, the microscope observations were performed between 2.5 and 5 min after the EGF addition. (**A**) Typical ultrafast (6 kHz) single fluorescent-molecule trajectories of EGFR, before and after EGF stimulation (*n* = 41 and 39 trajectories, respectively). (**B**) Distributions of the compartment sizes detected by the single-molecule diffusion of EGFR in the basal PM, before (**top**) and after (**bottom**) EGF stimulation (for the method, see the caption to Fig. 2 **C**). Arrowheads indicate the median values of 106 nm for both before and after stimulation. (**C**) Distributions of the EGFR residency times within a compartment determined by the TILD analysis, with the best-fit exponential curves, providing the dwell lifetimes (τ). Before (**top**) and after (**bottom**) EGF stimulation. Statistically significant difference between before and after stimulation with *P* = 0.044, using the log-rank test. (**D**) Distributions of *D*_MACRO_ for EGFR in the basal PM determined by the hop-diffusion fitting of the MSD-Δ*t* plot for each EGFR molecule (shaded histogram, before stimulation; green open histogram, after stimulation). Arrowheads indicate the median values. *D*_MACRO_ was reduced by a factor of 1.7 after stimulation.

The dwell lifetime of EGFR within a compartment was increased from 18 ± 1.2 ms before stimulation to 27 ± 1.5 ms during 2.5 − 5 min after stimulation (**Fig. 3** **C**), which reduced *D*_MACRO_ by ∼42% (**Fig. 3** **D**). Since EGFR is likely to form stable dimers upon stimulation, the slowed macroscopic diffusion or elongated dwell lifetime within a compartment might be induced by the lower hop probabilities of the larger diffusant; i.e., engaged EGFR dimers. This result is consistent with the “oligomerization-induced trapping” model proposed previously (Iino et al., 2001; Murakoshi et al., 2004; Heinemann et al., 2013). The longer confinement of engaged receptors within or near the compartment where the EGF signal was originally received will be beneficial for localizing the signal, which might in turn induce the local reorganization of the actin cytoskeleton required for membrane ruffling and chemotaxis.

### The sizes and inter-island distances of paxillin islands in the FA determined by live-cell ultrafast PALM

Using single-molecule imaging, the FA region was previously found to be quite accessible by non-FA proteins, as well as integrin and other FA molecules, allowing them to undergo diffusion there (Shibata et al., 2012, 2013; Rossier et al., 2012; Tsunoyama et al., 2018; Orré et al., 2021). These results demonstrated that the massive protein assembly (continent) and Venetian canal models would not be suitable for the FA protein organization in the FA (**Fig. 1** **D**). Before we further investigated the molecular dynamics in the FA region, we quantitatively observed the distribution of a prototypical FA protein, paxillin, in live cells using ultrafast PALM. Paxillin can bind to integrin and is reportedly located near the PM (Kanchanawong et al., 2010; Liu et al., 2015).

We performed ultrafast PALM observations with a data acquisition rate of 1 kHz for 10 s (10,000 frames), using the developed camera system and live T24 cells expressing mEos3.2 fused to human paxillin (**Fig. 4** **A top**). The average number of mEos3.2-paxillin molecules detected in the entire basal PM (using a PALM data acquisition frame size of 640 x 640 pixels) was approximately 300,000 copies/cell, when almost all of them were photobleached. Assuming that 60% of mEos3.2 is fluorescent (Baldering et al., 2019) and that 30% of mEos3.2 paxillin is recruited to the basal PM, then 1,650,000 copies of mEos3.2-paxillin would be expressed in a cell. Since 920,000 copies of zyxin and 810,000 copies of VASP are expressed in a T24 cell (Tsunoyama and Kusumi, private communication), it would be reasonable to assume that ∼800,000 copies of endogenous paxillin exist in a T24 cell. Therefore, the paxillin in the cells used for PALM would be overexpressed by a factor of ∼3.1.

**Figure 4.**
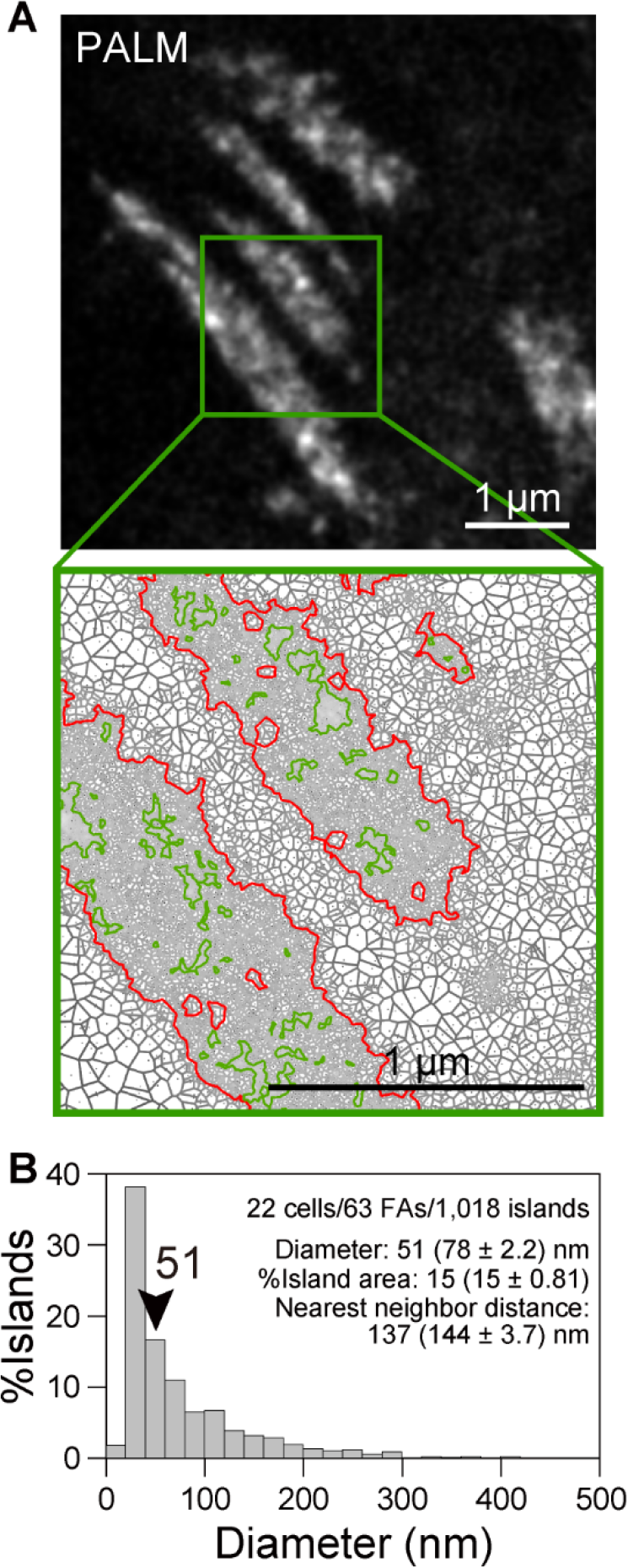
The FA is likely to be an archipelago of FA-protein islands (as detected with paxillin) of 51-nm median diameter dispersed in the compartmentalized fluid, with a median nearest-neighbor distance of 137 nm. (**A**) (**top**) Typical ultrafast PALM image of mEos3.2-paxillin expressed on the basal PM of a live T24 cell (data acquisition at 1 kHz for 10 s, with a total of 10,000 frames). (**bottom**) The Voronoï polygons’ expression of the image enclosed in the square in the top image. The contours of FAs (red) and FA-protein islands (green) are shown. (**B**) Distribution of the diameters of FA-protein islands, providing the median diameter of 51 nm. The nearest-neighbor distance was 137 nm (median). See **Materials and methods**, “Ultrahigh-speed imaging of single Halo-TfR and EGFR-Halo labeled with TMR and ultrafast live-cell PALM of mEos3.2-paxillin”.

The obtained images were similar to those reported previously (Shroff et al., 2007, 2008; Changede and Sheetz, 2017; Deschout et al., 2017; Orré et al., 2021). By applying the SR-Tesseler software (Levet et al., 2015) to the PALM images (**Fig. 4** **A bottom**), we obtained the diameter distribution of the paxillin islands (**Fig. 4** **B**). The median diameter was 51 nm and the median nearest-neighbor inter-island distance was 137 nm. The paxillin islands occupy 15 ± 0.81% of the FA area. These results are consistent with the finding that both non-FA and FA molecules enter the FA zone quite readily and undergo diffusion there, and that the fluid membrane occupies the majority of the FA region.

### TfR undergoes hop diffusion inside the FA

To observe the dynamics of membrane molecules in the fluid membrane part of the FA region, we observed the movements of the non-FA TM protein TfR inside the FA at an ultrafast frame rate of 6 kHz (0.167-ms resolution). As described, outside the FA in the basal PM, TfR undergoes hop diffusion (**Figs. 2 and 5**; **Movie S1 top**). However, we were struck to find that, when TfR molecules entered the FA zone (marked by mGFP-paxillin), they kept undergoing hop diffusion (**Fig. 5, middle and bottom**; **Movie S1 bottom**).

**Figure 5.**
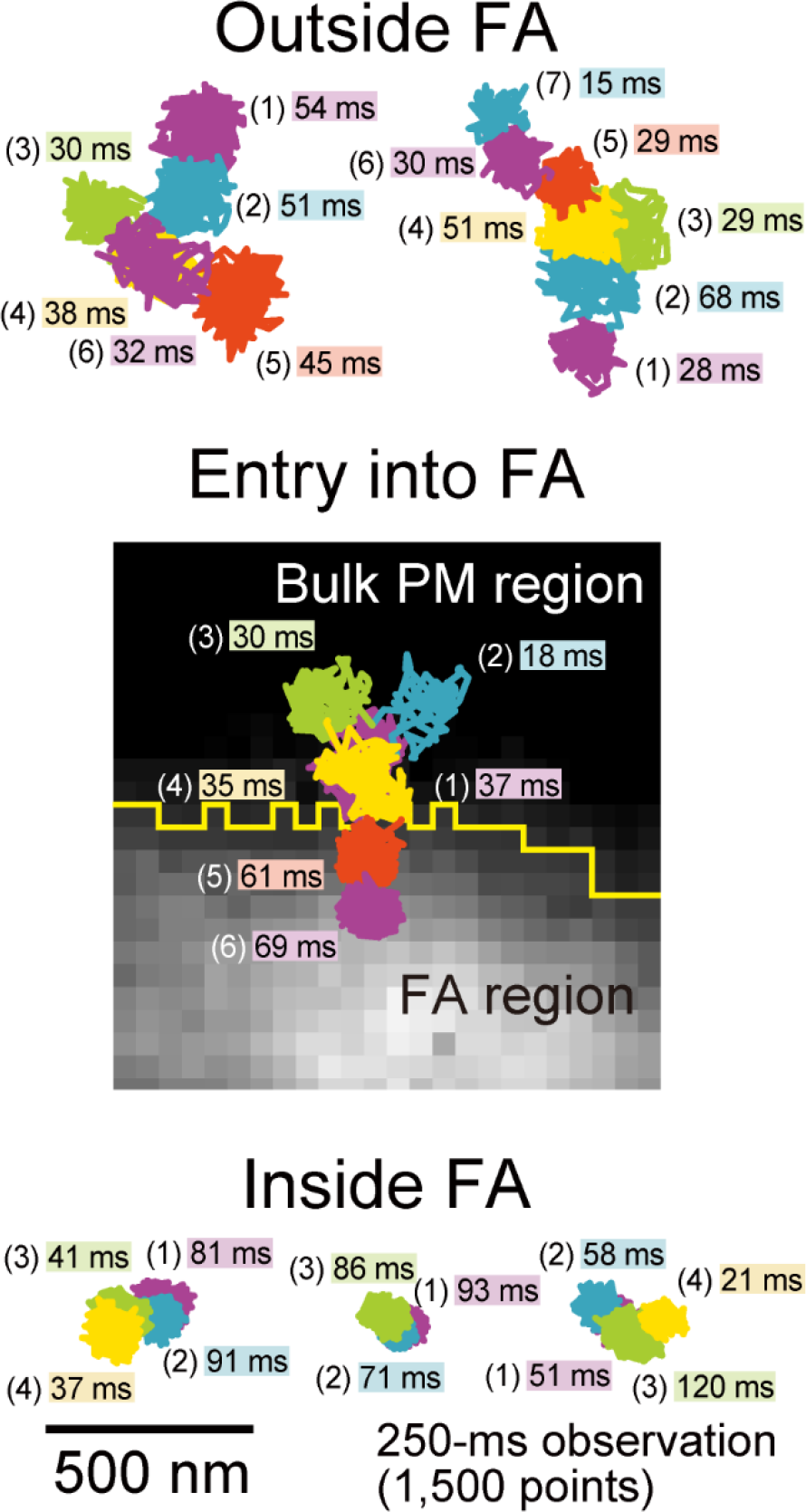
Typical ultrafast single fluorescent-molecule trajectories of TfR observed at 6 kHz, diffusing outside the FA region (top), entering the FA region from the bulk PM (middle), and diffusing inside the FA zone (bottom). In the middle figure, the background of the trajectory is the mGFP-paxillin image, and the yellow line shows the boundary of the FA region determined by binarization, using the minimum cross entropy thresholding.

A quantitative analysis showed that 85% of the TfR trajectories inside the FA zone were statistically classified into the suppressed diffusion mode (**Fig. 6** **A top**; **Table 1**). The hop-diffusion fitting of the MSD-*Δt* plot of each trajectory (see the caption to **Fig. S5** of the companion paper) revealed that the median compartment size was reduced to 74 nm (from 109 nm in the bulk basal PM, **Fig. 6** **B**). Namely, the compartment area size was reduced by a factor of 2.2. The dwell lifetime was increased to 36 ± 2.8 ms in the FA (from 24 ± 1.6 ms in the bulk basal PM, **Fig. 6** **C**). These results unequivocally demonstrated that the fluid membrane in the inter-island channels in the FA is compartmentalized. Due to the similarity to the hop diffusion in the bulk PM (Fujiwara et al., 2002, 2016; Murase et al., 2004), we propose that the compartmentalization would be induced by the meshwork of the actin-based membrane skeleton. However, attempts to directly observe the effects of actin depolymerizing drugs failed, because at the concentrations where their effects were detectable, the cells became round and some did not survive. Following the model of hop diffusion in actin-induced PM compartmentalization, the present result indicates that the actin-based membrane skeleton meshwork is finer in the FA and more stable. Stabilized actin meshwork, perhaps due to the smaller mesh size and/or the binding of the proteins enriched in the FA region, would allow the passage of membrane molecules across the compartment boundaries less frequently (see **Discussion**).

**Figure 6.**
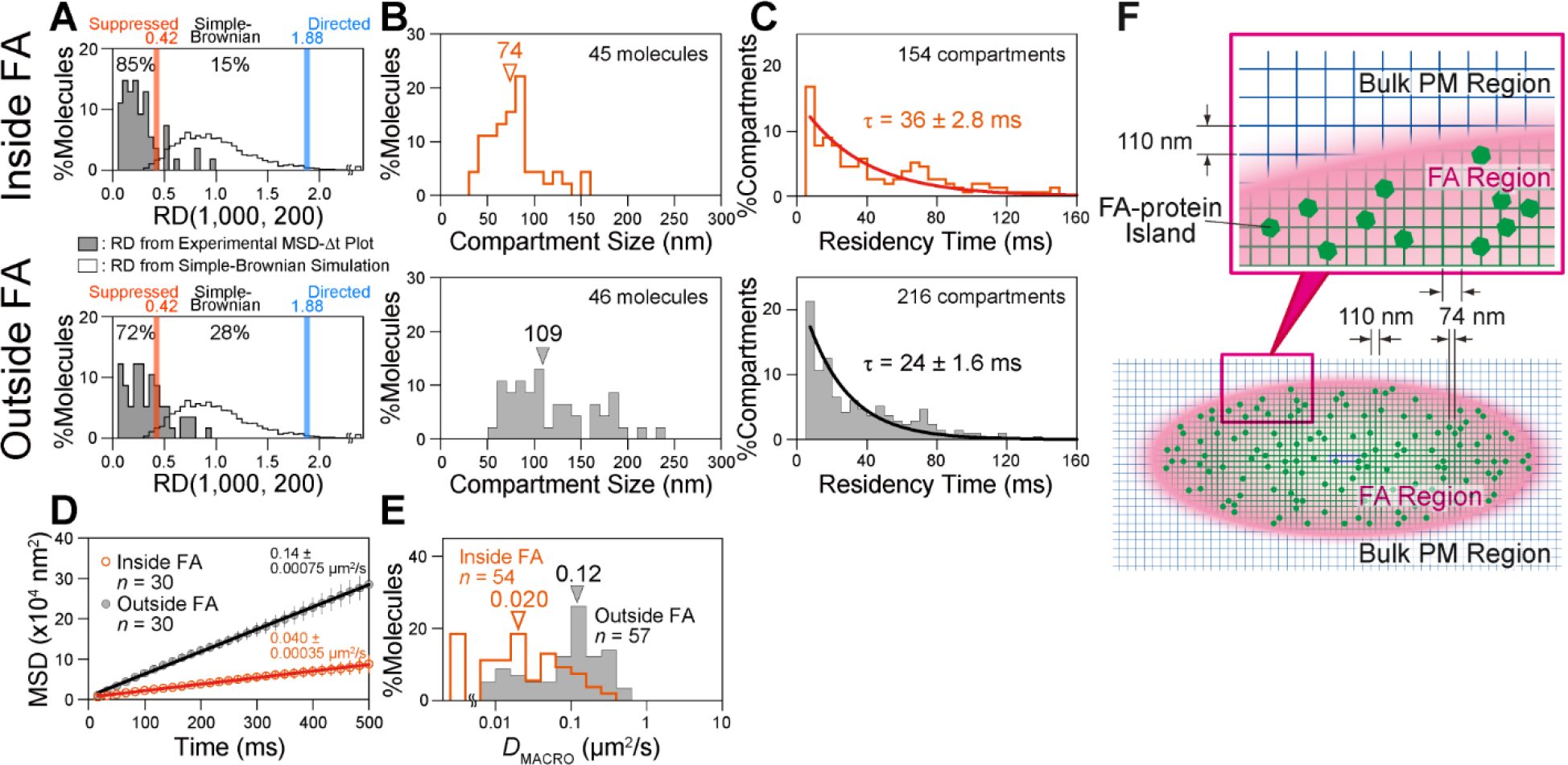
TfR undergoes hop diffusion within the FA region, as in the bulk basal PM, but with a smaller median compartment size of 74 nm (vs. 109 nm) and a longer dwell lifetime of 36 ms (vs. 24 ms). (**A**) Distributions of *𝑅𝐷*s for the trajectories of TfR diffusing inside (**top**) and outside (**bottom**) the FA region in the basal PM (shaded histograms; 54 and 57 molecules, respectively). The open histogram (the same for both inside and outside the FA) shows the *𝑅𝐷* distribution for simple-Brownian trajectories generated by Monte-Carlo simulations (5,000 trajectories) and is the same as that shown in Fig. 2 **B**. (**B**) Distributions of the compartment sizes determined by the hop-diffusion fitting of the MSD-Δ*t* plot of TfR for each trajectory, inside (**top**) and outside (**bottom**) the FA region in the basal PM. Arrowheads indicate the median values (Statistically significant difference with *P* = 1.7 x 10^-7^, using the Brunner-Munzel test). (**C**) Distributions of the TfR residency times within a compartment determined by the TILD analysis, with the best-fit exponential curves (providing the dwell lifetimes), inside (**top**) and outside (**bottom**) the FA region. Statistically significant difference between before and after stimulation with *P* = 4.9 x 10^-4^, using the log-rank test. (**D**) Ensemble-averaged MSD-Δ*t* plots for TfR obtained from the observations at 60-Hz (every 16.7 ms). At this frame rate, the hop behaviors are totally smeared out as seen in the linear plot here. The slope of the ensemble-averaged MSD-Δ*t* plot (divided by 4) provides *D*_500ms_, representing the long-term diffusion coefficient for molecules undergoing many hops across the compartments and being affected by other obstacles in the PM. (**E**) Distributions of *D*_MACRO_ for TfR determined by the hop-diffusion fitting of the MSD-Δ*t* plot for each TfR trajectory obtained at 6 kHz. Arrowheads indicate the median values (statistically significant difference with *P* = 9.3 x 10^-9^, using the Brunner-Munzel test). (**F**) Schematic model of the FA architecture. The FA-protein islands (paxillin islands) of 51 nm in median diameter (green hexagons) in the archipelago are dispersed with the median closest neighbor distance of 137 nm in the compartmentalized fluid (magenta region with the green lattice). The fluid membrane in the inter-island channels in the FA is partitioned into 74-nm compartments (110 nm in the bulk PM; **B**). The compartment boundaries are likely to be composed of the actin-based membrane-skeleton mesh. This mesh might be bound and stabilized by various FA proteins (green meshes due to the binding of FA proteins; outside the FA, the meshes are blue, showing that the properties of the actin meshes would be different).

### The compartmentalized fluid membrane in the FA is sparsely dotted with FA-protein islands

Further analysis of TfR diffusion was performed in the time scale of 500 ms, which is much longer than the time scale of the previous analyses of TfR diffusion in the time scale of ∼20 ms using the time resolution of 0.167 ms (6 kHz; see **Fig. 4** **A** in the companion paper). The observation frame rate was reduced to 60 Hz (time resolution of 16.7 ms). At this frame rate, hop diffusion is totally smeared out (hop diffusion features become undetectable) and the movement of TfR appears like simple-Brownian diffusion (consistent with most available data for molecular diffusion in the PM observed at video rate, 30 Hz, in the literature), but single molecules could be observed much longer (the number of frames in a trajectory is basically the same, but the duration in terms of real time is longer due to longer frame time of 16.7 ms, rather than 0.167 ms).

The ensemble-averaged MSD-Δ*t* plots (averaged over all obtained trajectories) are shown in **Fig. 6** **D**. The MSD-Δ*t* plots could be fitted by a straight line, indicating that at this slow observation frame rate, the TfR movements appear like simple-Brownian diffusion (for the detailed discussion, see **Fig. 5** **B** and the related text in the companion paper). The slope of the linear fitting provides the mean diffusion coefficient in the time scale of 500 ms (*D*_500ms_). The *D*_500ms_ values outside and inside the FA were 0.14 and 0.040 µm^2^/s, respectively. These values are quite comparable with the *D*_MACRO_ values obtained by the hop-diffusion fitting of the 6-kHz data in the time scale of 20 ms, which were 0.12 (0.14 ± 0.016) and 0.020 (0.046 ± 0.0081) µm^2^/s in median (mean ± SEM) values, outside and inside the FA respectively (**Fig. 6** **E**; **Table 1**). Since *D*_MACRO_ values obtained from the 6-kHz data in the time scale of 20 ms would mostly represent the occurrences of hops once and a few times in a trajectory, the similar values of *D*_MACRO_ (median = 0.020 µm^2^/s; mean = 0.046 µm^2^/s) and *D*_500ms_ (0.040 µm^2^/s) suggest that the long-term diffusion for 500 ms is determined by the compartmentalization of the fluid membrane, both inside and outside the FA. During 500 ms, inside the FA, TfR will diffuse over the distances of 0.28 µm (0.08 µm^2^ in the 2D area), which would be ∼15 compartments in 2D (using *D*_500ms_ = 0.040 µm^2^/s inside the FA and the median square compartment size of 74 nm). Namely, the TfR diffusion in the 1D space scale of 0.28 µm or 2D scale of ∼15 compartments inside the FA is rather dominated by the compartmentalization of the fluid membrane, rather than the diffusion-barrier-like effect of paxillin islands (and also the FA-protein islands; the obstacle effect of paxillin/FA-islands for TfR diffusion was undetectable).

Therefore, the TfR diffusion measurements clearly indicates that the paxillin/FA-islands are dispersed quite sparsely in the FA so that their areal occupancy in the FA region would be less than 20% (Holcman et al., 2011; Saxton, 1982, 2010). Although the direct comparisons would be difficult, these results are generally consistent with our PALM observations of the paxillin islands: the paxillin islands occupy 15% of the FA area, their median diameter, 51 nm, is smaller than the median compartment size (74 nm), and the median nearest-neighbor inter-island distance was 137 nm (**Fig. 4** **B**). This result further suggests that the FA-islands that do not contain paxillin would be quite limited in number and/or much smaller.

Based on these observations, we propose an FA architectural model, which we call the “*archipelago of the FA-protein islands in the compartmentalized fluid*” (**Fig. 6** **F**). In this model, membrane molecules enter the FA domain more or less freely (Shibata et al., 2012, 2013; Rossier et al., 2012; Tsunoyama et al., 2018; Orré et al., 2021), undergoing hop diffusion. The rapid diffusion of integrins and other FA proteins and their preassembled oligomers (Hoffmann et al., 2014) inside the FA domain would facilitate their rapid exchanges with those located in the bulk domain, allowing the simultaneous formation and disintegration of the FA-islands (including paxillin islands) everywhere in the FA region (the formation and disintegration do not have to occur from the outer boundary of the FA), and thus accelerating the FA formation-disintegration. The FA-protein islands might be bound to the actin-based membrane skeleton mesh for their immobilization, which would facilitate the maintenance of the FA zone identity in the bulk basal PM.

### Integrin becomes temporarily immobilized at FA-protein islands

We next performed simultaneous live-cell PALM of mEos3.2-paxillin for visualizing the FA-protein islands and single-molecule tracking of integrin β3 (integrin β3’s N-terminus fused to the ACP-tag protein, labeled with SeTau647; <1,000 copies expressed on the cell surface, thus avoiding overexpression) at 250 Hz (**Fig. 7, A-D**; **Movie S3**). The distributions of the paxillin pixel signal intensities in the FA zone obtained from the PALM images (normalized by the median value) are shown in **Fig. 7** **D-top**. Meanwhile, each time the integrin β3 molecule exhibited immobilization in the FA (some immobilizations lasted for much longer than the photobleaching lifetime of 270 frames = 1.1 s at 250 Hz, see Tsunoyama et al., 2018), we measured the paxillin pixel signal intensity at the center of the immobilized position (Simson et al., 1995) (**Fig. 7, B and C**). After observing many temporary immobilizations of single integrin β3 molecules, we obtained the histogram of the paxillin signal intensities at the places where the temporary immobilization of integrin β3 occurred (**Fig. 7** **D-bottom**). More than two-thirds of the temporary immobilization events occurred where the paxillin intensity was greater than its median value, indicating that integrin β3 molecules are preferentially immobilized at places where the paxillin densities are high; namely, on the paxillin islands, dynamically linking them to the extracellular matrix. The linkage of the paxillin island to the extracellular matrix would be accomplished by many integrin molecules, which are dynamically exchanging all the time. The protein compositions of the FA-protein islands could vary, and some FA-protein islands may contain only a few paxillin molecules, which we might have missed. Therefore, we propose that integrin molecules dynamically and transiently associate with FA-protein islands and the extracellular matrix. The dynamic linkage would facilitate the FA formation and disintegration (Shroff et al., 2008).

**Figure 7.**
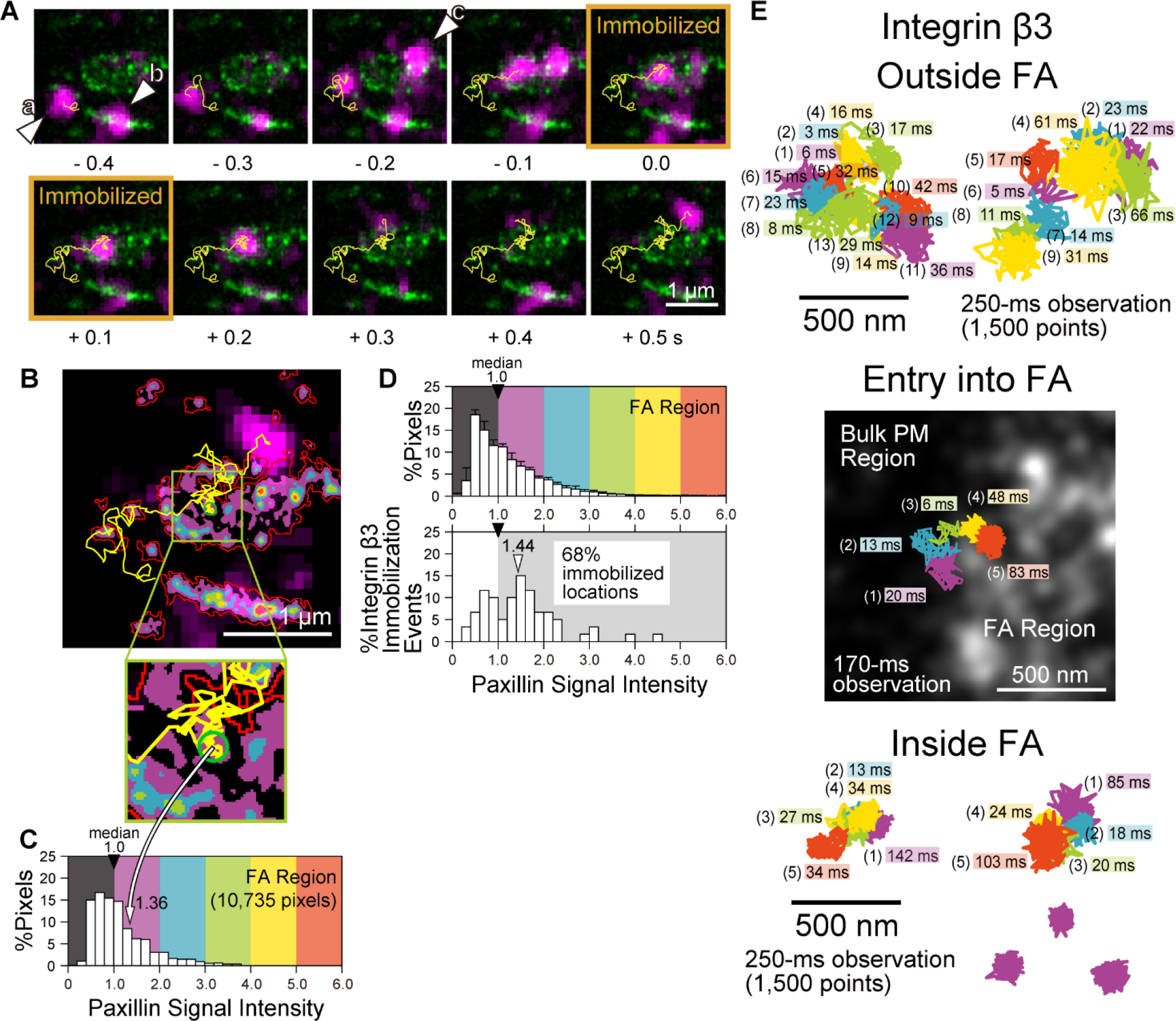
Integrin β3 molecules are intermittently and transiently immobilized at paxillin islands. (**A**) Typical results of simultaneous live-cell PALM of mEos3.2-paxillin, visualizing the paxillin islands (green spots; the same image is used for all of the panels) and single-molecule tracking of SeTau647-ACP-integrin β3 (recorded at a 4-ms resolution; large magenta spots; images shown every 0.1 s or 25 frames). Time 0 is when one of the ACP-integrin β3 molecules (arrowhead **a**, with trajectories, among the three molecules **a**, **b**, and **c**) started exhibiting immobilization, which lasted for 0.19 s (orange frames). Since the sizes of the green spots of the FA-protein islands are determined by the localization errors, they are much smaller than the magenta spots of single integrin molecules, with sizes determined by the diffraction limit. The signal intensities of the magenta spots at times 0.3 and 0.4 s are lower, probably due to the stochastic fluctuation of emitted photon numbers and blinks (off periods) shorter than 4 ms. (**B**) The thermographic PALM image of mEos3.2-paxillin shown in **A**, together with the FA boundaries determined by the minimum cross entropy thresholding (red contours). The immobilized area, as previously defined (Simson et al., 1995), for molecule **a** (see the yellow trajectory) is shown by a green circle with a 104-nm diameter (see the central part of the green square and the expanded figure below). The thermographic color scale is shown in **C**. (**C**) Distribution of mEos3.2-paxillin signal intensities in the FA regions shown in **B** (normalized by the median signal intensity), shown with the thermographic color scale. (**D**) (**top**) Normalized distribution of mEos3.2-paxillin signal intensities, averaged over 60 FAs. (**bottom**) Distribution of normalized mEos3.2-paxillin signal intensities (per pixel) at the centers of the immobilization circles of single integrin molecules (for an example, see the green circle in **B**; *n* = 60 events), indicating that integrin molecules are preferentially immobilized on the FA-protein islands (see the main text). The arrowhead indicates the median value. (**E**) Typical ultrafast single fluorescent-molecule trajectories of integrin β3 observed at 6 kHz, diffusing outside the FA region (**top**), entering the FA region from the bulk PM (**middle**), and diffusing and becoming temporarily immobilized inside the FA zone (**bottom**). The middle figure shows a typical result of simultaneous observations of SeTau647-Halo-integrin β3 entering the FA region from the bulk PM using ultrafast (6 kHz) single-molecule tracking (colored trajectory) and the paxillin islands visualized by ultrafast live-cell PALM of mEos3.2-paxillin (1 kHz, 10-s integration). Consistent with the FA model of the archipelago architecture of FA-protein islands in compartmentalized fluid, integrin molecules continued to exhibit hop diffusion when they entered the FA region, outside the paxillin islands.

Finally, the frame rate for observing integrin β3 (integrin β3’s N-terminus fused to the Halo-tag protein, labeled with SeTau647) was increased from 250 Hz to 6 kHz, and the integrin molecules entering the FA zone from the bulk membrane were observed. Typical results are shown in **Fig. 7** **E** and **Movie S4**. Consistent with the FA model of the archipelago architecture of FA-protein islands in the compartmentalized fluid, integrin β3 molecules exhibited hop diffusion in the bulk PM, continued the hop diffusion after they entered the FA zone, and then became immobilized. More systematic research results of integrin β3 hop diffusion will be published elsewhere.

## Discussion

We report the following four findings in this paper, which were obtained by using the ultrafast single-molecule imaging and ultrafast PALM described in the companion paper. First, we addressed the question that previously arose often about whether the bulk basal PM is significantly different from the bulk apical PM. We found that the basal PM is compartmentalized in virtually the same manner as the apical PM (**Fig. 2**). Not only the compartment sizes but also the dwell lifetimes found for Cy3-DOPE and TfR in the basal PM outside the FA were virtually the same as those found in the apical PM. Indeed, we were quite surprised to find these. We propose that the compartmentalization of the bulk PM is induced by the actin-based membrane skeleton meshes attached to the PM cytoplasmic surface (picket-fence model, **Fig. 1** **B**; addressing the question outside the FA in **Fig. 1** **C**). It follows then that the actin-based membrane skeleton has very similar structures and dynamics on the cytoplasmic surfaces of the apical and basal PMs.

Second, using EGFR, we confirmed the compartmentalization of the basal PM (**Fig. 3**). The dwell lifetime of EGFR within a compartment was 18 ms, which is shorter than that of TfR (24 ms). This is consistent with the previous findings that EGFR exists as both monomers and dimers in dynamic equilibrium before EGF stimulation (Chung et al., 2010; Low-Nam et al., 2011), whereas TfR is a constitutive dimer (and both are single-pass transmembrane proteins), because monomers would move across the compartment boundaries more readily than dimers (Fujiwara et al., 2016). Upon EGF ligation, the dwell lifetime of EGFR within a compartment was increased to 27 ms (from 18 ms before stimulation) without any changes in the compartment size. This is consistent with the fact that ligated EGFR forms stabilized dimers and greater oligomers (Chung et al., 2010; Low-Nam et al., 2011; Huang et al., 2016). The present results, obtained before and after EGF stimulation, support the “oligomerization-induced trapping” model. Dimers and oligomers would undergo intercompartmental hop movements less frequently, because dimers and oligomers would require more space than monomers to move between two transmembrane picket proteins and/or two severed actin filaments in the actin-induced membrane-skeleton meshes. If membrane-bound proteins are floating in a simple two-dimensional fluid, then the diffusion coefficients of oligomers, say 10-mers, would not be appreciably smaller than monomers (Saffman and Delbrück, 1975). Therefore, oligomerization-induced trapping in the actin-induced partitioned PM is critical for understanding the molecular behaviors in the PM.

Third, we examined the superhigh speed single-molecule dynamics of both the non-FA protein TfR and the FA protein β3-integrin in the FA region, as a specialized domain in the basal PM. Unexpectedly, we found that both TfR and β3-integrin undergo hop diffusion in the FA domain, suggesting that the bulk liquid membrane part in the FA region is also compartmentalized (**Figs. 5, 6, 7E**). Quantitative analyses showed that the median compartment size in the FA (74 nm) is smaller than that in the bulk PM (106 - 109 nm, found using TfR, Cy3-DOPE, and EGFR), and that the TfR’s dwell lifetime within a compartment in the FA is 36 ms, whereas that in the bulk PM is 24 ms (addressing the question in the FA in **Fig. 1** **C**). The smaller compartment sizes and longer molecular dwell lifetimes within the compartment in the FA, as compared with the bulk PM, explain the slower macroscopic diffusion in the FA by factors of 2 - 3 relative to that in the bulk PM (**Fig. 6, D and E**). These results strongly argue against the massive protein assembly (continent) model and the Venetian canal model (**Fig. 1** **D, left and middle**), and indicate that, if the FA-protein islands’ archipelago model (**Fig. 1** **D, right**) is correct, then the islands’ areal occupancy in the FA region would be less than 20% (Holcman et al., 2011; Saxton, 1982, 2010).

We believe that the compartmentalization of the FA region is induced by the actin-based membrane skeleton meshes attached to the PM cytoplasmic surface (picket-fence model, **Fig. 1** **B**), due to the similarities in the hop diffusion in the bulk basal PM (and thus the bulk apical PM). As described, we could not perform direct examinations using actin-depolymerizing and stabilizing reagents, because the high reagent concentrations required for modulating the hop diffusion in the FA region greatly affected the overall cell shape and viability. Assuming the validity of the actin-induced compartmentalization model, how could the dwell lifetime of TfR within a compartment be prolonged (how could the hop frequency across the compartment boundary be reduced) in the FA region as compared with the bulk basal PM? If the properties of the compartment boundaries; i.e., the actin fence and the transmembrane protein pickets, were the same as those in the bulk basal PM, then since the compartment size is smaller in the FA region than in the bulk PM (medians of 74 nm vs. 106 - 109 nm ; **Fig. 6, B and F**), the hop frequency would increase by a factor of more than 2 ([108/74]^2^) due to the increased frequency of TfR arrival at the compartment boundaries. However, the hop frequency was reduced by ∼30%, from every 36 ms to every 24 ms (**Fig. 6** **C**). This result suggests that the actin-based membrane skeleton mesh in the FA region is more stable and/or the average densities of the protein pickets anchored to and aligned along the actin mesh are higher.

We propose that both of these phenomena occur in the FA. In addition to the paxillin clusters, representing many FA-protein islands that were visible by PALM, there would also be various smaller FA-protein complexes bound to actin filaments in the FA region that were not clearly detectable as paxillin clusters by PALM. These actin-bound FA-proteins could decrease the thermal structural fluctuation of actin filaments and the frequencies of temporary severing of actin filaments, which would induce the brief collapse of compartment boundaries. The actin-bound FA-proteins, possibly including integrins, would also provide the effects of steric hindrance as well as the long-range fluid dynamic friction on molecules arriving at the compartment boundary regions, reducing their crossing probabilities.

Fourth, by performing simultaneous live-cell PALM of paxillin and single integrin β3 molecule imaging-tracking, we found that integrin molecules diffusing in the fluid membrane region in the FA preferentially undergo temporary immobilizations, primarily at the paxillin islands (**Fig. 7** **D**). We propose that the temporary binding of β3 integrin to the paxillin islands would be responsible for FA binding to the extracellular matrix and its regulation. Multiple integrin molecules would exist in a paxillin island and undergo dynamic exchanges with integrin molecules outside the paxillin islands inside the FA region. Namely, integrin molecules are continually recruited to the paxillin islands, and each recruited integrin molecule temporarily links the paxillin island to the extracellular matrix, and then leaves the paxillin island. Subsequently, other integrin molecules assume the responsibility for the linkage, again temporarily. Such a mechanism would be important for cell locomotion. During cell locomotion, tension on the FA would vary greatly, and depending on its strength, the dynamic linkage-exchange mechanism would allow the paxillin island to change the number of integrin molecules engaged in binding to the extracellular matrix on the island. In this sense, it would be interesting to examine the differences between β3 and β1 integrin molecules in terms of their immobilizations at the paxillin islands and their dynamics, because β1 and β3 integrins play distinct roles in cell adhesion and migration (Roca-Cusachs et al., 2009; Rossier et al., 2012; Schiller et al., 2013): β1 integrin might be important at the initial stages of the FA changes (growth/disintegration), whereas β3 integrin might be more important at the maintenance stages of the FA (Tsunoyama et al., 2018).

Approximately one-third of the β3 integrin’s temporary immobilization events took place outside the paxillin islands (**Fig. 7** **D**). This might be due to the binding of integrin to protein islands that lack paxillin and/or to smaller FA-protein clusters bound to the actin-based membrane skeleton meshes. In either case, the FA-protein clusters/islands must contain molecules with integrin binding sites and molecules with actin binding sites. These preformed clusters might mature into larger FA-protein islands (Hoffmann et al., 2014), although they may or may not contain paxillin.

We note that the binding of integrin to other FA molecules and FA-protein clusters floating in the fluid membrane-part of the FA region, and probably confined in the actin-based compartments, will not be detected as immobilizations here. Since the single-molecule localization precisions of TMR (bound to transmembrane proteins) in the basal PM is on the order of 35 nm, and the median compartment size in the FA is 74 nm, the confined integrin molecules that are not binding to actin meshes would not appear to be immobilized.

Further investigations of the localizations of paxillin outside the paxillin islands and the spatial distributions of other FA proteins in the FA are important future tasks. In addition, determining the FA protein distributions in the z (height) direction in each island would be informative. The means by which the actin-based membrane skeleton meshes in the FA are linked to the stress fibers must be clarified. As such, there are still many unresolved questions. However, we emphasize the importance of our conclusion: the FA largely consists of the fluid membrane partitioned into 74-nm compartments by actin-based picket fences, and the paxillin/FA-protein islands of 51-nm in the median diameter are dotted with the median nearest neighbor distance of 137 nm on the actin meshes, occupying 15% of the FA area, as represented by the archipelago model of the FA-protein islands in the compartmentalized fluid (**Fig. 6** **F**).

## Materials and methods

### Cell culture

Human T24 epithelial cells (the ECV304 cell line used in the previous research, which turned out to be a sub-clone of T24; Dirks et al., 1999; Murase et al., 2004) were grown in Ham’s F12 medium (Sigma-Aldrich) supplemented with 10% fetal bovine serum (Sigma-Aldrich). Cells were cultured on 12-mm diameter glass-bottom dishes (IWAKI), and single-molecule observations were performed two days after inoculation. For culturing cells expressing mGFP-paxillin or mEos3.2-paxillin, the glass surface was coated with fibronectin by an incubation with 5 µg/ml fibronectin (Sigma-Aldrich) in phosphate-buffered saline (PBS) (pH 7.4) at 37°C for 3 h.

### Fluorescence labeling of TfR and integrin β3 for observations of diffusion in the FA region: cDNA construction, expression, and labeling in T24 cells

The cDNA encoding human TfR (GenBank: M11507.1) fused to the Halo-tag protein at the TfR’s N-terminus (Halo-TfR) was generated by replacing the cDNA encoding the EGFP protein, in the EGFP-TfR plasmid, with that of the Halo-tag protein (Promega), with the insertion of a 45-base linker (15 amino acids, with the sequence SGGGG x3) between Halo and TfR. The cDNA encoding human paxillin isoform alpha (NCBI reference sequence: NM_002859.3; cloned from the human WI-38 cell line) fused to mGFP at the N-terminus was subcloned into the EBV-based episomal vector, pOsTet15T3 (gift from Y. Miwa, University of Tsukuba, Japan), which bears the tetracycline-regulated expression units, the transactivator (rtTA2-M2), and the TetO sequence (a Tet-on vector) (Shibata et al., 2012, 2013). To avoid perturbing the FA structure by overexpressing mGFP-paxillin, doxycycline induction was not employed, but leaky expression was used. However, due caution is required when interpreting how effectively mGFP-paxillin imitates the behaviors of native paxillin. The cDNA encoding mEos3.2-paxillin was generated by replacing the cDNA encoding mGFP in the mGFP-paxillin plasmid with that encoding mEos3.2 (Zhang et al., 2012) (generated by making 3 point mutations, I102N, H158E, and Y189A in the Addgene mEos2 plasmid #20341 [http://n2t.net/addgene:20341; RRID:Addgene_20341], a gift from L. Looger, Janelia Research Campus; McKinney et al., 2009). The cDNA encoding human integrin β3 (NCBI reference sequence: NM_000212.2; a gift from J. C. Jones, Northwestern University; Tsuruta et al., 2002) tagged with ACP (ACP-integrin β3) or Halo (Halo-integrin β3) was subcloned into the pOsTet15T3 vector. For ACP-integrin β3, the CD8 signal peptide before ACP and a 15-base linker (5 amino acids, with the sequence SGGGG) between ACP and integrin β3 were employed, whereas for Halo-integrin β3, the IL6 signal peptide before Halo and a 45-base linker (15 amino acids, with the sequence SGGGG x3) between Halo and integrin β3 were used. The leaky expression without doxycycline induction was selected to avoid overexpression. All of the newly generated constructs were confirmed by DNA sequencing.

T24 cells were transfected with the cDNA encoding mGFP-paxillin, using the Lipofectamine LTX and Plus reagents (Invitrogen) and with other cDNAs by using Nucleofector 2b (Lonza), following the manufacturers’ recommendations. To covalently link TMR to Halo-TfR, T24 cells coexpressing Halo-TfR and mGFP-paxillin were incubated in Hanks’ balanced salt solution, buffered with 2 mM TES (Dojindo) at pH 7.4 (HT medium), containing 10 nM TMR-conjugated Halo-ligand (Promega) at 37°C for 1 h, and then washed three times with HT medium. The remaining unbound ligand in the cytoplasm was removed by incubating the cells in HT medium for 30 min, and then washing the cells three times with HT medium. To form the covalent complex between SeTau647 (SETA BioMedicals) and ACP-integrin β3, T24 cells coexpressing ACP-integrin β3 and mEos3.2-paxillin were incubated in cell culture medium (Ham’s F12 medium supplemented with 10% fetal bovine serum) containing 100 nM SeTau647-CoA (Shinsei Chemical Company), 1 µM ACP synthase (New England Biolabs), and 10 mM MgCl_2_ for 30 min at 37°C, and then washed three times with HT medium. To form the covalent complex between SeTau647 and Halo-integrin β3, T24 cells coexpressing Halo-integrin β3 and mEos3.2-paxillin were incubated in cell culture medium containing 100 nM SeTau647-conjugated Halo-ligand (Shinsei Chemical Company) at 37°C for 1 h, and then washed three times with HT medium. We previously found that under these conditions, >90% of ACP and Halo could be complexed with their fluorescent ligands (Morise et al., 2019). ACP-integrin β3 was used for observing temporary immobilizations at a frame rate of 250 Hz, due to the smaller size of the ACP-tag protein as compared with the Halo-tag protein, for a possibly smaller effect on the integrin β3 binding to FA-protein islands. Halo-integrin β3 was used for observing hop diffusion at 6 kHz, due to the better stability of the SeTau647 dye bound to the Halo-tag protein at higher laser intensities, as compared with the dye bound to the ACP-tag protein.

### EGFR-Halo expression in T24 cells and its TMR labeling

The cDNA encoding human EGFR (GenBank: X00663.1) tagged with Halo (EGFR-Halo) was generated by replacing the cDNA encoding the YFP protein, in the EGFR-YFP plasmid (a gift from I. R. Nabi, University of British Columbia, Canada), with that of the Halo 7-tag protein (Promega), with the insertion of a 63-base linker (21 amino acids, with the sequence RVPRDPSGGGGSGGGGSGGGG) between EGFR and Halo.

T24 cells were transfected with the cDNA encoding EGFR-Halo, using a Nucleofector 2b device, and then starved in serum-free Ham’s F12 medium for ∼16 h. The covalent complex between TMR and EGFR-Halo was formed in the same way as in the labeling of Halo-TfR. For EGF stimulation, HT medium containing 20 nM human EGF (Sigma-Aldrich) was added to the cells at a final concentration of 10 nM (1 ml of 20 nM EGF in HT medium was added to the glass-bottom dish containing 1 ml HT medium). The measurements were performed before and during 2.5 - 5 min after the EGF addition.

### Ultrahigh-speed imaging of single Halo-TfR and EGFR-Halo labeled with TMR and ultrafast live-cell PALM of mEos3.2-paxillin

Each individual Halo-TfR and EGFR-Halo on the basal PM was observed at 37 ± 1°C, using the TIR illumination mode of a home-built objective lens-type TIRF microscope (based on an Olympus IX70 inverted microscope), which was modified and optimized for the camera system developed in the companion paper. A 532-nm laser (Millennia Pro D2S-W, 2W, Spectra-Physics) for exciting TMR was attenuated with neutral density filters, circularly polarized, and then steered into the edge of a high numerical aperture (NA) oil immersion objective lens (UAPON 150XOTIRF, NA = 1.45, Olympus), focused on the back-focal plane of the objective lens.

For ultrafast live-cell PALM of mEos3.2-paxillin, as well as the detection and characterization of paxillin islands using Voronoï-based segmentation of the PALM images (**Fig. 4**), see the **Materials and methods**, “Ultrafast live-cell PALM of caveolin-1-mEos3.2 and mEos3.2-paxillin (Figs. 6-8)” and “Detection and characterization of FA-protein islands using Voronoï-based segmentation of the PALM images (Fig. 8)” of the companion paper. The diameter of each paxillin island was determined by the principal-component analysis (PCA) provided in the SR-Tesseler software, and the mean nearest neighbor distance of the FA-protein islands within each FA region was calculated as 1/(2*ρ*^1/2^), where *ρ* represents the number density of islands in each FA. The results are shown in **Fig. 4, A** and **B**.

**Simultaneous live-cell PALM of mEos3.2-paxillin, to visualize the FA-protein islands, and high-speed imaging of SeTau647-ACP (or Halo)-integrin β3, to track their single-molecule movements in the FA archipelago (Fig. 7)** Simultaneous data acquisitions of ultrafast live-cell PALM of mEos3.2-paxillin and ultrafast single-molecule tracking of ACP-integrin β3 or Halo-integrin β3 labeled with SeTau647 were performed on the basal PM at 37 ± 1°C. Ultrafast PALM was performed as described in the **Materials and methods** of the companion paper.

Single-molecule imaging of SeTau647 bound to integrin β3 was performed (simultaneously with the ultrafast live-cell PALM data acquisition) at a frame rate of 250 Hz for visualizing the integrin β3 binding to the FA-protein islands (using ACP-integrin β3) or 6 kHz for visualizing the integrin β3 hop diffusion inside the FA (using Halo-integrin β3), using a 660-nm laser (Ventus, 750 mW, Laser Quantum) at 0.84 µW/µm^2^ or 8.4 µW/µm^2^, respectively. In the excitation arm, a multi-band mirror (ZT405/488/561/647rpc, Chroma) was employed. The leakage of the intense 561-nm excitation laser beam into the emission arm was blocked by a notch filter (NF03-561E, Semrock) placed right before the entrance of the detection arm. The fluorescence images of mEos3.2 and SeTau647 were split by a dichroic mirror (ZT647rdc, Chroma) into two detection arms with bandpass filters of 572-642 nm for mEos3.2 (FF01-607/70, Semrock) and 672-800 nm for SeTau647 (FF01-736/128, Semrock). Each detection arm was equipped with the developed ultrafast camera system, and the images were projected onto the photocathode of the camera system (more specifically, the photocathode of the image intensifier). The images from the two cameras were spatially corrected and superimposed with sub-pixel precisions, as described previously (Koyama-Honda et al., 2005).

## Data availability

Data supporting the findings of this study are available from the corresponding author upon reasonable request.

## Code availability

The code is available from the corresponding author upon reasonable request.

## Online supplemental material

**Fig. S1** shows that the effect of photo response non-uniformity (PRNU) of the developed camera system on the single-molecule localization precision is minimal. **Video 1** shows single TfR (Halo-TMR) molecules diffusing outside and inside the FA regions in the basal PM of T24 cells, observed at 6 kHz (every 0.167 ms). **Video 2** shows single EGFR (Halo-TMR) molecules diffusing in the basal PM before and 2.5 min after the addition of 10 nM EGF, observed at 6 kHz (every 0.167 ms). **Video 3** shows a single integrin β3 molecule (ACP-SeTau647) intermittently binding to FA-protein islands defined by the live-cell PALM of mEos3.2-paxillin, observed at 250 Hz (every 4 ms). **Video 4** shows an integrin β3 molecule (Halo-SeTau647) exhibiting hop diffusion in the bulk basal PM, and continuing to exhibit hop diffusion after it entered the FA zone visualized by the live-cell PALM of mEos3.2-paxillin, observed at 6 kHz (every 0.167 ms).

## Acknowledgements

We thank Profs. Y. Miwa of the University of Tsukuba, J. C. Jones of Northwestern University, I. R. Nabi of the University of British Columbia, and L. Looger of the Janelia Research Campus, for their kind gifts of the pOsTet15T3 vector, the cDNA encoding human integrin β3, the human EGFR-YFP plasmid, and the mEos2 plasmid, respectively. We also thank K. Hanaka for his enthusiastic encouragement of this research, M. Yahara and A. Chadda for constructing various cDNAs, J. Kondo-Fujiwara and K. Kanemasa for preparing the figures, and the members of the Kusumi laboratory for helpful discussions. We are grateful to Prof. Ken Jacobson of the University of North Carolina and Prof. Kathalina Gaus of the University of New South Wales Sydney for critical reading of the manuscript and constructive comments. This work was supported in part by Grants-in-Aid for Scientific Research from the Japan Society for the Promotion of Science (JSPS) (Kiban B to T.K.F. [16H04775, 20H02585], Kiban B to K.G.N.S. [18H02401], and Kiban S and A to A.K. [16H06386 and 21H04772, respectively]), a Grant-in-Aid for Challenging Research (Exploratory) from JSPS to T.K.F. (18K19001), a Grant-in-Aid for Innovative Areas from the Ministry of Education, Culture, Sports, Science and Technology of Japan (MEXT) to K.G.N.S. (18H04671), a Japan Science and Technology Agency (JST) grant in the program of the Core Research for Evolutional Science and Technology (CREST) in the field of “Biodynamics” to A.K. and that in the field of “Extracellular Fine Particles” to K.G.N.S. (JPMJCR18H2), and a JST grant in the program of the Development of Advanced Measurements and Analysis Systems to A.K. and T.K.F. WPI-iCeMS of Kyoto University is supported by the World Premiere Research Center Initiative (WPI) of MEXT.

## Author contributions

T.K.F. and A.K. conceived and formulated the project. T.K.F., S.T., Y.N., K.I., and A.K. developed the ultrahigh-speed camera system, T.K.F., K.I., T.A.T., and A.K. developed an ultrafast single-molecule tracking station based on the newly developed camera system, and T.K.F., T.K., T.A.T., and A.K. tested the camera system on the developed station. T.K.F., A.C.E.S., T.A.T., L.H.C., K.G.N.S., and A.K. designed the biological experiments and participated in discussions. T.K.F. performed virtually all of the ultrahigh-speed single-molecule imaging-tracking experiments, the ultrafast PALM experiments, and the data analysis. T.K.F. and K.P.R. developed the compartment detection algorithm. Z.K., based on discussions with T.K.F., derived the equation describing the MSD-Δ*t* plot for particles undergoing hop diffusion and developed the theory for the distribution of the residency times within a compartment for particles undergoing hop diffusion. T.K.F., K.P.R., Z.K., and A.K. evaluated the data. T.K.F., Z.K., and A.K. wrote the manuscript, and all authors discussed the results and participated in revising the manuscript.

## Supplementary Figure

**Figure S1.**
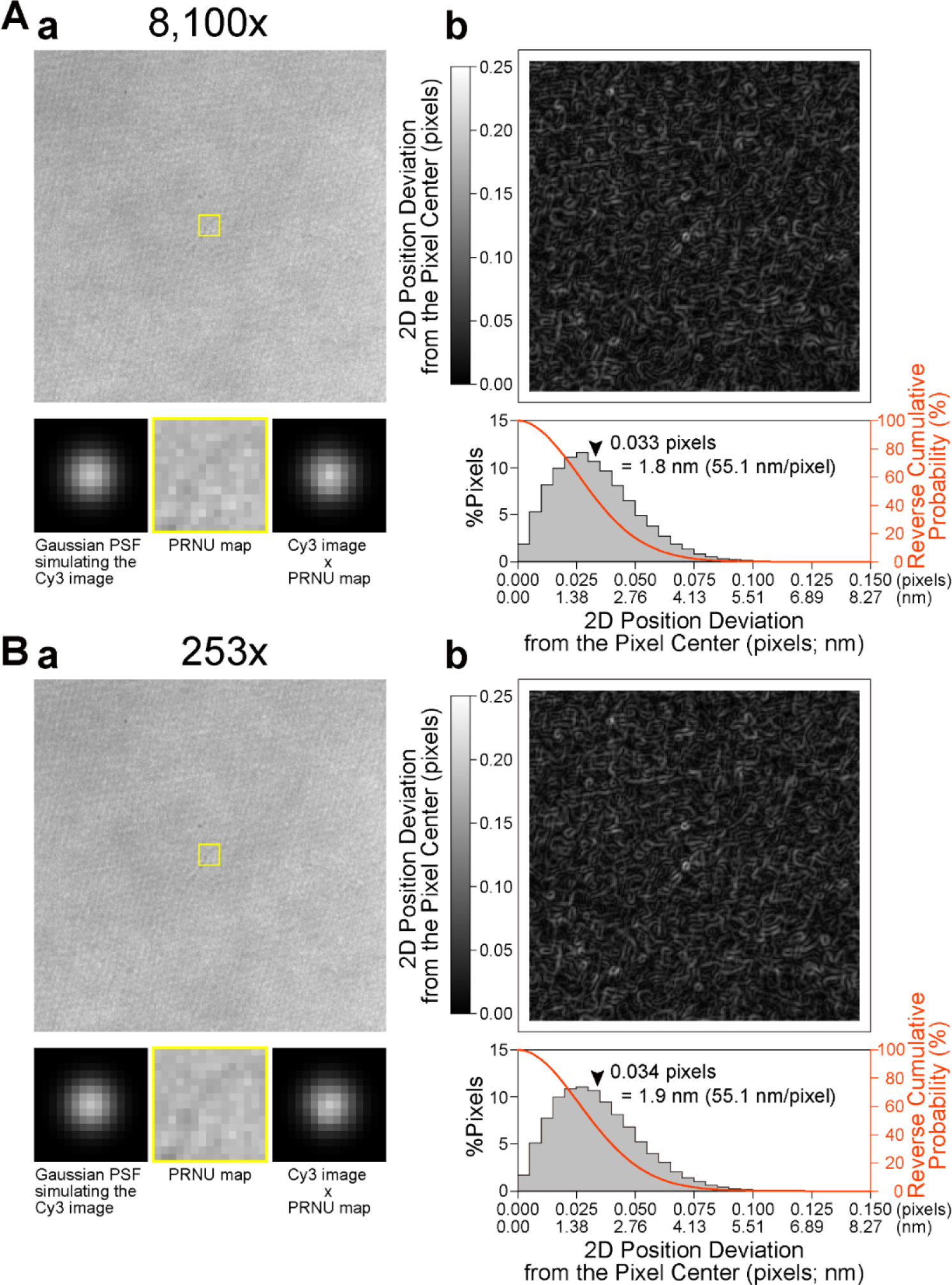
The photo response non-uniformity (PRNU) of the developed camera system scarcely affects the single-molecule localization precision, examined at 10 kHz using two image intensifier amplifications of 8,100x (A) and 253x (B). (**a**) (**top**) 256 x 256-pixel images representing PRNU of the developed camera system, obtained by averaging images over 40,000 consecutive frames recorded at 10 kHz under uniform illumination, so that the mean pixel intensity counts became 515 ± 34 (**A**) and 513 ± 32 (**B**) (SD for 256 x 256 pixels), which are about half of the maximum intensity count of 10 bits. The uniform illumination was generated by Köhler illumination, using the halogen lamp of the microscope and a 572-642-nm bandpass filter (FF01-607/70, Semrock). (**bottom**) Modulation of the image of a single Cy3 molecule by PRNU, evaluated by calculations. (**left**) The Cy3 image was approximated by an ideal two-dimensional Gaussian point spread function (PSF) in the 15 x 15-pixel region, based on an experimentally determined standard deviation of 2.2 pixels for 50 Cy3 molecules immobilized on the glass, obtained by the TIR illumination at 79 µW/µm^2^ and a peak intensity count of 511 (half of the maximum intensity count of 10 bits). In the actual imaging experiments, we employed 55.1 nm/pixel: 2.2 pixels = 123 nm. The PSF peak was placed at the center of the 15 x 15-pixel region. (**middle**) The 15 x 15-pixel yellow regions shown in the **top** images are magnified. (**right**) Images on the **left** and **middle** were multiplied pixel-by-pixel and normalized, generating the PRNU-modulated images of a single Cy3 molecule. (**b**) The effect of PRNU on the single-molecule localization precision is quite limited. (**top**) Maps of the 2D position deviation from the pixel center (coded on the grey scale). These maps were generated by moving the 15 x 15-pixel image of an ideal Gaussian PSF simulating the Cy3 image (**a, bottom-left**), scanning over the 256 x 256-pixel PRNU images (**a, top**) pixel-by-pixel, and calculating the 2D position deviation at every position in the scan. (**bottom**) Distributions of the 2D position deviations showing that the mean deviations are 0.033 and 0.034 pixels (= 1.8 nm and 1.9 nm at 55.1 nm/pixel; *n* = 57,600 pixels; arrowheads) for (**A**) and (**B**), respectively, which are comparable to the typical PRNU effect found with the EM-CCD camera (Pertsinidis et al., 2010). Furthermore, the reverse cumulative distributions (shown in red) indicate that 95% of the 2D position deviations are within the range of 0.066 pixels (= 3.6 nm) and 0.068 pixels (= 3.7 nm) for (**A**) and (B), respectively. These results suggest that the effect of PRNU on the single-molecule localization precision is limited and virtually the same at amplifications of 8,100x (**A**) and 253x (**B**).

## Captions for Videos 1 to 4

### Video 1

Single TfR (Halo-TMR) molecules diffusing in the *basal* PM of T24 cells, observed at 6 kHz (0.167-ms frame time) (observation period of 1,500 frames = 250 ms; Replay, 50x-slowed from real time). Movies on the top (bottom) show the movement of a single TfR molecule outside (inside) the FA regions (see **Fig. 2** and its caption). Movies on the left (larger view-fields): green areas, FA regions marked by mGFP-paxillin; magenta spots, single TfR molecules. The regions in the yellow squares are enlarged in the movies on the right, showing single-molecule trajectories outside (top) or inside (right) the FA zone. Within the FA region, the compartment size is smaller by a factor of 2 in area and the dwell lifetime within a compartment is ∼50% longer, as compared with those outside the FA region.

### Video 2

Single EGFR (Halo-TMR) molecules diffusing in the intact *basal* PM before (left) and 2.5 min after (right) the addition of 10 nM EGF, observed at 6 kHz (every 0.167 ms) (observation period of 1,500 frames = 250 ms; Replay, 50x-slowed from real time) (see **Fig. 5** and its caption). The statistical analysis method developed here classified both trajectories into the suppressed-diffusion mode. Engaged EGFR molecules hopped less frequently, as compared with non-ligated EGFR, probably due to dimerization/oligomerization (oligomerization-induced trapping).

### Video 3

A single integrin β3 molecule (magenta spot with a yellow trajectory) intermittently binding to FA-protein islands (green regions) defined by the live-cell PALM (see **Fig. 7** **A** and its caption). The live-cell PALM of mEos3.2-paxillin was simultaneously performed with single-molecule imaging of ACP(SeTau647 label)-integrin β3 molecules at 250 Hz (every 4 ms; observation period = 1 s [250 frames]; Replay, 8.3x-slowed from real time). The arrowhead indicates the immobilization event.

### Video 4

An integrin β3 molecule exhibiting hop diffusion in the bulk basal PM, and continuing to exhibit hop diffusion after it entered the FA zone. The live-cell PALM of mEos3.2-paxillin (green regions = FA-protein islands; still image due to a 10-s data acquisition time = 1-ms integration time/frame x 10,000 frames) was simultaneously performed with ultrafast single-molecule imaging of a single Halo(SeTau647 label)-integrin β3 molecule at 6 kHz (magenta spot with a color-coded trajectory; every 0.167 ms; observation period = 170 ms [1,024 frames]; Replay, 50x-slowed from real time) (see **Fig. 7** **E** and its caption).

## References

Baldering T.N., M.S. Dietz, K. Gatterdam, C. Karathanasis, R. Wieneke, R. Tampé, and M. Heilemann. 2019. Synthetic and genetic dimers as quantification ruler for single-molecule counting with PALM. Mol. Biol. Cell. 30:1369–1376. doi: 10.1091/mbc.E18-10-0661. https://doi.org/10.1091/mbc.E18-10-0661

Bays, J.L., and K.A. DeMali. 2017. Vinculin in cell-cell and cell-matrix adhesions. Cell Mol. Life Sci. 74:2999–3009. https://doi.org/10.1007/s00018-017-2511-3

Calderwood, D.A., I.D. Campbell, and D.R. Critchley. 2013. Talins and kindlins: partners in integrin-mediated adhesion. Nat. Rev. Mol. Cell Biol. 14:503–517. https://doi.org/10.1038/nrm3624

Changede, R., and M. Sheetz. 2017. Integrin and cadherin clusters: A robust way to organize adhesions for cell mechanics. Bioessays. 39:1–12. https://doi.org/10.1002/bies.201600123

Chung, I., R. Akita, R. Vandlen, D. Toomre, J. Schlessinger, and I. Mellman. 2010. Spatial control of EGF receptor activation by reversible dimerization on living cells. Nature 464:783-787. https://doi.org/10.1038/nature08827

Deschout, H., I. Platzman, D. Sage, L. Feletti, J.P. Spatz, and A. Radenovic. 2017. Investigating focal adhesion substructures by localization microscopy. Biophys. J. 113:2508–2518. https://doi.org/10.1016/j.bpj.2017.09.032

Dirks, W.G., R.A.F. MacLeod, and H.G. Drexler. 1999. ECV304 (endothelial) is really T24 (bladder carcinoma): Cell line cross-contamination at source. In Vitro Cell Dev. Biol. Anim. 35:558–559. https://doi.org/10.1007/s11626-999-0091-8

Fujiwara, T., K. Ritchie, H. Murakoshi, K. Jacobson, and A. Kusumi. 2002. Phospholipids undergo hop diffusion in compartmentalized cell membrane. J. Cell Biol. 157:1071–1081. https://doi.org/10.1083/jcb.200202050

Fujiwara, T.K., K. Iwasawa, Z. Kalay, T.A. Tsunoyama, Y. Watanabe, Y.M. Umemura, H. Murakoshi, K.G. Suzuki, Y.L. Nemoto, N. Morone, and A. Kusumi. 2016. Confined diffusion of transmembrane proteins and lipids induced by the same actin meshwork lining the plasma membrane. Mol. Biol. Cell. 27:1101–1119. https://doi.org/10.1091/mbc.E15-04-0186

Gardel, M.L., I.C. Schneider, Y. Aratyn-Schaus, and C.M. Waterman. 2010. Mechanical integration of actin and adhesion dynamics in cell migration. Annu. Rev. Cell Dev. Biol. 26:315–333. https://doi.org/10.1146/annurev.cellbio.011209.122036

Heinemann, F., S. Vogel, and P. Schwille. 2013. Lateral membrane diffusion modulated by a minimal actin cortex. Biophys. J. 104:1465–1475. https://doi.org/10.1016/j.bpj.2013.02.042

Hoffmann, J.E., Y. Fermin, R.L. Stricker, K. Ickstadt, and E. Zamir. 2014. Symmetric exchange of multi-protein building blocks between stationary focal adhesions and the cytosol. Elife. 3:e02257. https://doi.org/10.7554/eLife.02257

Holcman, D., N. Hoze, and Z. Schuss. 2011. Narrow escape through a funnel and effective diffusion on a crowded membrane. Phys. Rev. E Stat. Nonlin. Soft Matter Phys. 84:021906. https://doi.org/10.1103/PhysRevE.84.021906

Huang, Y., S. Bharill, D. Karandur, S.M. Peterson, M. Marita, X. Shi, M.J. Kaliszewski, A.W. Smith, E.Y. Isacoff, and J. Kuriyan. 2016. Molecular basis for multimerization in the activation of the epidermal growth factor receptor. Elife. 5:e14017. https://doi.org/10.7554/eLife.14107

Humphries, J.D., M.R. Chastney, J.A. Askari, and M.J. Humphries. 2019. Signal transduction via integrin adhesion complexes. Curr. Opin. Cell Biol. 56:14–21. https://doi.org/10.1016/j.ceb.2018.08.004

Iino, R., I. Koyama, and A. Kusumi. 2001. Single molecule imaging of green fluorescent proteins in living cells: E-cadherin forms oligomers on the free cell surface. Biophys. J. 80:2667–2677. https://doi.org/10.1016/S0006-3495(01)76236-4

Kanchanawong, P., G. Shtengel, A.M. Pasapera, E.B. Ramko, M.W. Davidson, H.F. Hess, and C.M. Waterman. 2010. Nanoscale architecture of integrin-based cell adhesions. Nature. 468:580–584. https://doi.org/10.1038/nature09621

Koseska A., and P.I. Bastiaens. 2020. Processing temporal growth factor patterns by an epidermal growth factor receptor network dynamically established in space. Annu. Rev. Cell Dev. Biol. 36:359–383. https://doi 10.1146/annurev-cellbio-013020-103810

Koyama-Honda, I., K. Ritchie, T. Fujiwara, R. Iino, H. Murakoshi, R. Kasai, and A. Kusumi. 2005. Fluorescence imaging for monitoring the colocalization of two single molecules in living cells. Biophys. J. 88:2126–2136. https://doi.org/10.1529/biophysj.104.048967

Krause, M., E.W. Dent, J.E. Bear, J.J. Loureiro, and F.B. Gertler. 2003. Ena/VASP proteins: regulators of the actin cytoskeleton and cell migration. Annu. Rev. Cell Dev. Biol. 19:541–564. https://doi.org/10.1146/annurev.cellbio.19.050103.103356

Kusumi, A., Y. Sako, and M. Yamamoto. 1993. Confined lateral diffusion of membrane receptors as studied by single particle tracking (nanovid microscopy). Effects of calcium-induced differentiation in cultured epithelial cells. Biophys. J. 65:2021–2040.

Levet, F., E. Hosy, A. Kechkar, C. Butler, A. Beghin, D. Choquet, and J.B. Sibarita. 2015. SR-Tesseler: a method to segment and quantify localization-based super-resolution microscopy data. Nat. Methods. 12:1065–1071. https://doi.org/10.1016/S0006-3495(93)81253-0

Liu, J., Y. Wang, W.I. Goh, H. Goh, M.A. Baird, S. Ruehland, S. Teo, N. Bate, D.R. Critchley, M.W. Davidson, and P. Kanchanawong. 2015. Talin determines the nanoscale architecture of focal adhesions. Proc. Natl. Acad. Sci. U.S.A. 112:E4864–E4873. https://doi.org/10.1073/pnas.1512025112

Liu, S., D.A. Calderwood, and M.H. Ginsberg. 2000. Integrin cytoplasmic domain-binding proteins. J. Cell Sci. 113 (Pt 20):3563–3571. https://doi.org/10.1242/jcs.113.20.3563

Low-Nam, S.T., K.A. Lidke, P.J. Cutler, R.C. Roovers, P.M. van Bergen en Henegouwen, B.S. Wilson, and D.S. Lidke. 2011. ErbB1 dimerization is promoted by domain co-confinement and stabilized by ligand binding. Nat. Struct. Mol. Biol. 18:1244–1249. https://doi.org/10.1038/nsmb.2135

López-Colomé, A.M., I. Lee-Rivera, R. Benavides-Hidalgo, and E. López. 2017. Paxillin: a crossroad in pathological cell migration. J. Hematol. Oncol. 10:50. https://doi.org/10.1186/s13045-017-0418-y

McKinney, S.A., C.S. Murphy, K.L. Hazelwood, M.W. Davidson, and L.L. Looger. 2009. A bright and photostable photoconvertible fluorescent protein. Nat. Methods. 6:131–133. https://doi.org/10.1038/nmeth.1296

Morise, J., K. Suzuki, A. Kitagawa, Y. Wakazono, K. Takamiya, T. Tsunoyama, Y. Nemoto, H. Takematsu, A. Kusumi, and S. Oka. 2019. AMPA receptors in the synapse turnover by monomer diffusion. Nat. Commun. 10:5245. https://doi.org/10.1038/s41467-019-13229-8

Morone, N., T. Fujiwara, K. Murase, R.S. Kasai, H. Ike, S. Yuasa, J. Usukura, and A. Kusumi. 2006. Three-dimensional reconstruction of the membrane skeleton at the plasma membrane interface by electron tomography. J. Cell Biol. 174:851–862. https://doi.org/10.1083/jcb.200606007

Murakoshi, H., R. Iino, T. Kobayashi, T. Fujiwara, C. Ohshima, A. Yoshimura, and A. Kusumi. 2004. Single-molecule imaging analysis of Ras activation in living cells. Proc. Natl. Acad. Sci. U.S.A. 101:7317–7322. https://doi.org/10.1073/pnas.0401354101

Murase, K., T. Fujiwara, Y. Umemura, K. Suzuki, R. Iino, H. Yamashita, M. Saito, H. Murakoshi, K. Ritchie, and A. Kusumi. 2004. Ultrafine membrane compartments for molecular diffusion as revealed by single molecule techniques. Biophys. J. 86:4075–4093. https://doi.org/10.1529/biophysj.103.035717

Nayal, A., D.J. Webb, C.M. Brown, E.M. Schaefer, M. Vicente-Manzanares, and A.R. Horwitz. 2006. Paxillin phosphorylation at Ser273 localizes a GIT1-PIX-PAK complex and regulates adhesion and protrusion dynamics. J. Cell Biol. 173:587–589. https://doi.org/10.1083/jcb.200509075

Orré, T., A. Joly, Z. Karatas, B. Kastberger, C. Cabriel, R.T. Böttcher, S. Lévêque-Fort, J.B. Sibarita, R. Fässler, B. Wehrle-Haller, O. Rossier, and G. Giannone. 2021. Molecular motion and tridimensional nanoscale localization of kindlin control integrin activation in focal adhesions. Nat. Commun. 12:3104. https://doi.org/10.1038/s41467-021-23372-w

Parsons, J.T., A.R. Horwitz, and M.A. Schwartz. 2010. Cell adhesion: integrating cytoskeletal dynamics and cellular tension. Nat. Rev. Mol. Cell Biol. 11:633–643. https://doi.org/10.1038/nrm2957

Patla, I., T. Volberg, N. Elad, V. Hirschfeld-Warneken, C. Grashoff, R. Fässler, J.P. Spatz, B. Geiger, and O. Medalia. 2010. Dissecting the molecular architecture of integrin adhesion sites by cryo-electron tomography. Nat. Cell Biol. 12:909–915. https://doi.org/10.1038/ncb2095

Reynolds A.R., C. Tischer, P.J. Verveer, O. Rocks, and P.I. Bastiaens. 2003. EGFR activation coupled to inhibition of tyrosine phosphatases causes lateral signal propagation. Nat. Cell Biol. 5:447–453. https://doi 10.1038/ncb981

Roca-Cusachs, P., N.C. Gauthier, A. del Rio, and M.P. Sheetz. 2009. Clustering of α_5_β_1_ integrins determines adhesion strength whereas α_V_β_3_ and talin enable mechanotransduction. Proc. Natl. Acad. Sci. U.S.A. 106:16245–16250. https://doi.org/10.1073/pnas.0902818106

Rossier, O., V. Octeau, J.B. Sibarita, C. Leduc, B. Tessier, D. Nair, V. Gatterdam, O. Destaing, C. Albiges-Rizo, R. Tampe, L. Cognet, D. Choquet, B. Lounis, and G. Giannone. 2012. Integrins β1 and β3 exhibit distinct dynamic nanoscale organizations inside focal adhesions. Nat. Cell Biol. 14:1231–1231. https://doi.org/10.1038/Ncb2620

Saffman, P.G., and M. Delbrück. 1975. Brownian motion in biological membranes. Proc. Natl. Acad. Sci. U.S.A. 72:3111–3113. https://doi.org/10.1073/pnas.72.8.3111

Saxton, M.J. 1982. Lateral diffusion in an archipelago. Effects of impermeable patches on diffusion in a cell membrane. Biophys. J. 39:165–173. https://doi.org/10.1016/S0006-3495(82)84504-9

Saxton, M.J. 2010. Two-dimensional continuum percolation threshold for diffusing particles of nonzero radius. Biophys. J. 99:1490–1499. https://doi.org/10.1016/j.bpj.2010.06.033

Schiller, H.B., M.R. Hermann, J. Polleux, T. Vignaud, S. Zanivan, C.C. Friedel, Z. Sun, A. Raducanu, K.E. Gottschalk, M. Théry, M. Mann, and R. Fässler. 2013. β_1_- and α_v_- class integrins cooperate to regulate myosin II during rigidity sensing of fibronectin-based microenvironments. Nat. Cell Biol. 15:625–636. https://doi.org/10.1038/ncb2747

Shibata, A.C., T.K. Fujiwara, L. Chen, K.G. Suzuki, Y. Ishikawa, Y.L. Nemoto, Y. Miwa, Z. Kalay, R. Chadda, K. Naruse, and A. Kusumi. 2012. Archipelago architecture of the focal adhesion: membrane molecules freely enter and exit from the focal adhesion zone. Cytoskeleton (Hoboken). 69:380–392. https://doi.org/10.1002/cm.21032

Shibata, A.C., L.H. Chen, R. Nagai, F. Ishidate, R. Chadda, Y. Miwa, K. Naruse, Y.M. Shirai, T.K. Fujiwara, and A. Kusumi. 2013. Rac1 recruitment to the archipelago structure of the focal adhesion through the fluid membrane as revealed by single-molecule analysis. Cytoskeleton (Hoboken). 70:161–177. https://doi.org/10.1002/cm.21097

Shroff, H., C.G. Galbraith, J.A. Galbraith, and E. Betzig. 2008. Live-cell photoactivated localization microscopy of nanoscale adhesion dynamics. Nat. Methods. 5:417–423. https://doi.org/10.1038/nmeth.1202

Simson, R., E.D. Sheets, and K. Jacobson. 1995. Detection of temporary lateral confinement of membrane proteins using single-particle tracking analysis. Biophys. J. 69:989–993. https://doi.org/10.1016/s0006-3495(95)79972-6

Sjöblom, B., A. Salmazo, and K. Djinović-Carugo. 2008. Alpha-actinin structure and regulation. Cell Mol. Life Sci. 65:2688–2701. https://doi.org/10.1007/s00018-008-8080-8

Spiess, M., P. Hernandez-Varas, A. Oddone, H. Olofsson, H. Blom, D. Waithe, J.G. Lock, M. Lakadamyali, and S. Strömblad. 2018. Active and inactive β1 integrins segregate into distinct nanoclusters in focal adhesions. J. Cell Biol. 217:1929–1940. https://doi.org/10.1083/jcb.201707075

Stutchbury, B., P. Atherton, R. Tsang, D.Y. Wang, and C. Ballestrem. 2017. Distinct focal adhesion protein modules control different aspects of mechanotransduction. J. Cell Sci. 130:1612–1624. https://doi.org/10.1242/jcs.195362

Sulzmaier, F.J., C. Jean, and D.D. Schlaepfer. 2014. FAK in cancer: mechanistic findings and clinical applications. Nat. Rev. Cancer. 14:598–610. https://doi.org/10.1038/nrc3792

Suzuki, K., K. Ritchie, E. Kajikawa, T. Fujiwara, and A. Kusumi. 2005. Rapid hop diffusion of a G-protein-coupled receptor in the plasma membrane as revealed by single-molecule techniques. Biophys. J. 88:3659–3680. https://doi.org/10.1529/biophysj.104.048538

Tsunoyama, T.A., Y. Watanabe, J. Goto, K. Naito, R.S. Kasai, K.G.N. Suzuki, T.K. Fujiwara, and A. Kusumi. 2018. Super-long single-molecule tracking reveals dynamic-anchorage-induced integrin function. Nat. Chem. Biol. 14:497–506. https://doi.org/10.1038/s41589-018-0032-5

Tsuruta, D., M. Gonzales, S. Hopkinson, C. Otey, S. Khuon, R. Goldman, and J. Jones. 2002. Microfilament-dependent movement of the β3 integrin subunit within focal contacts of endothelial cells. FASEB J. 16:866–868. https://doi.org/10.1096/fj.01-0878fje

Verveer P.J., F.S. Wouters, A.R. Reynolds, and P.I. Bastiaens. 2000. Quantitative imaging of lateral ErbB1 receptor signal propagation in the plasma membrane. Science 290:1567–1570. https://doi:10.1126/science.290.5496.1567

Xia, S., and P. Kanchanawong. 2017. Nanoscale mechanobiology of cell adhesions. Semin. Cell Dev. Biol. 71:53–67. https://doi.org/10.1016/j.semcdb.2017.07.029

Yamada, K.M., and M. Sixt. 2019. Mechanisms of 3D cell migration. Nat. Rev. Mol. Cell Biol. 20:738–752. https://doi.org/10.1038/s41580-019-0172-9

Yang, J., L. Zhu, H. Zhang, J. Hirbawi, K. Fukuda, P. Dwivedi, J. Liu, T. Byzova, E.F. Plow, J. Wu, and J. Qin. 2014. Conformational activation of talin by RIAM triggers integrin-mediated cell adhesion. Nat. Commun. 5:5880. https://doi.org/10.1038/ncomms6880

Zhang, M., H. Chang, Y. Zhang, J. Yu, L. Wu, W. Ji, J. Chen, B. Liu, J. Lu, Y. Liu, J. Zhang, P. Xu, and T. Xu. 2012. Rational design of true monomeric and bright photoactivatable fluorescent proteins. Nat. Methods. 9:727–729. https://doi.org/10.1038/nmeth.2021

## Reference

Pertsinidis, A., Y. Zhang, and S. Chu. 2010. Subnanometre single-molecule localization, registration and distance measurements. Nature. 466:647–651. https://doi.org/10.1038/nature09163

